# The spectrum of diversity of nucleotide-binding leucine-rich repeat (NLR) genes in citrus and its relatives

**DOI:** 10.1101/2025.09.22.677818

**Authors:** Emmanuel Avila de Dios, Shree P. Thapa, Taylor R. Beaulieu, Tania Toruño, Daniil M. Prigozhin, Gitta Coaker, Danelle K. Seymour

**Author notes:** These authors contributed equally (Emmanuel Avila de Dios, Shree P. Thapa, Taylor R. Beaulieu).

## Abstract

Genomic clusters of immune genes, including those encoding nucleotide-binding leucine-rich repeat (NLR) proteins, are a model for exploring the dynamics of genomic regions in flux. Rapid sequence evolution of immune genes, including NLRs, and variation in their gene content, may enable long-lived plants, which lack adaptive immune systems, to keep pace with the fast evolution of pathogens. To explore the patterns and processes shaping the evolution of NLR gene content in a genus of long-lived tree species, we unified the annotation of NLR genes across 11 accessions (or 15 haplotypes) from the genus *Citrus* and its relatives, including three new diploid genome assemblies. A majority of NLRs were arranged in genomic clusters composed of paralogous genes, typically from a single gene family. Even larger clusters, with 10 or more NLRs, were limited to genes derived from one or few gene families. These patterns suggested that genomic clustering of NLRs arose through local expansion of phylogenetically related NLRs, but the mechanistic processes driving these patterns are not clear. Local gene duplication can be mediated by multiple processes, including transposon-mediated gene capture and subsequent proliferation, and non-allelic repair of double stranded breaks, including unequal recombination. Examples of retrotransposon-mediated duplication of NLRs were identified, but these were not sufficient to explain massive regional expansions. Signatures of unequal recombination are challenging to identify. Focusing on recent lineage-specific sequence duplications, at least one case of unequal recombination was identified, supporting a role for unequal recombination in shaping genomic variation in these regions.

## Introduction

Plant pathogens rapidly adapt to new environments and host immune systems must keep pace in order to thrive. Unlike animals, plants lack an adaptive immune system, and many have puzzled over how plants, with much longer generation times, are able to combat the pressure from fast evolving pathogens (J. D. G. Jones and Dangl 2006). Plants sense and respond to pathogen ingress using surface localized and intracellular immune receptors, with integrated cross-talk between both (Dodds and Rathjen 2010; Yuan et al. 2021). The first line of defense consists of a series of extracellular immune receptors that can sense conserved molecular signatures associated with pathogens at the cell surface. These extracellular receptors are typically receptor protein (RP) or receptor kinases (RK) (Dodds, Chen, and Outram 2024). The second layer of defense is mounted by intracellular immune receptors known as nucleotide-binding leucine-rich repeat (NLR) proteins that recognize pathogen effectors injected into the plant cell. NLRs are continually challenged to recognize and respond to a spate of new pathogen effector proteins and some NLR gene sequences have signatures of rapid evolution (Prigozhin and Krasileva 2021; Van de Weyer et al. 2019).

Eukaryotic genes are not usually arranged and ordered by function along chromosomes (J. M. Lee and Sonnhammer 2003). In plant genomes, NLRs are one of the few exceptions to this trend, with a majority of characterized NLRs residing in complex genomic clusters at specific chromosomal locations (Zhou et al. 2004; Shao et al. 2014; R. R. Q. Lee and Chae 2020; Jiao and Schneeberger 2020; Woudstra et al. 2024; Q. Li, Jiang, and Shao 2021). Many have proposed that the clustering of NLR gene sequences promotes their rapid evolution, but evidence linking genomic clustering of NLRs with their pace of evolution is limited. Classical genetic studies provided early evidence of the existence of NLR clusters and led to hypotheses regarding the possible mechanisms shaping genetic variation in these clusters (Whitham et al. 1994; Bent et al. 1994; Mindrinos et al. 1994). In 1998, Michelmore and Meyers proposed that NLRs evolve through recurrent cycles of gene duplication and gene loss and that these “birth and death” cycles enabled plants to rapidly evolve NLRs with new specificities to keep up with the fast pace of pathogen evolution (Michelmore and Meyers 1998). This is still the working model for NLR evolution today. Early on it was thought that patterns of NLR clustering and massive presence and absence variation in NLR content was driven by unequal recombination (Hulbert 1997), which could quickly generate new combinations of NLR gene content. Michelmore and Meyers argued that non-homologous recombination would result in homogenization of NLR content and associated sequences and instead favored a combination of mutational processes leading to sequence divergence in clustered regions (Michelmore and Meyers 1998). Indeed, mispairing during mitosis and meiosis and subsequent ectopic/unequal recombination do occur (J. G. Jelesko et al. 1999), but their contribution to shaping genetic variation at NLR clusters is still unclear (Barragan and Weigel 2021).

Gene duplication can be a source of evolutionary novelty, including for NLRs (Flagel and Wendel 2009). Genes are duplicated through multiple processes including whole-genome duplication, transposon-mediated gene capture and subsequent proliferation, and non-allelic repair of double stranded breaks, including ectopic and unequal recombination. These latter processes of non-homologous recombination are favored for their potential to explain regional genomic clustering of NLR genes along chromosomes. Examples of each have been associated with clustered NLR sequences (D. Leister et al. 1998; Hulbert 1997), but their relative contribution to the evolution of this class of immune receptors is still unclear. Ectopic recombination occurs when a double stranded break is repaired with DNA from a distant genomic location. Phylogenetically related NLRs that are physically dispersed on different chromosomes is a signature of ectopic recombination (Dario Leister 2004; McDowell and Simon 2006). There is a clear example of a novel cluster of NLR genes in common bean arising from an initial ectopic recombination event where the donor NLR was located on a completely separate chromosome (David et al. 2009). While ectopic recombination can explain movement of NLR paralogs throughout the genome, unequal recombination is invoked to account for regional expansion of NLR genes, giving rise to their genomic clustering. Unequal recombination occurs when double stranded breaks are repaired with neighboring non-allelic sequences and resolution of these events leads to expansion or contraction of the intervening sequences (J. G. Jelesko et al. 1999). Tandem duplication of NLR genes is common (Jupe et al. 2012; Guo et al. 2011; G. Andolfo et al. 2013) and consistent with a role for unequal recombination in these regions (D. Leister et al. 1998). Painstaking characterization of recombination breakpoints at specific NLR clusters, are further evidence that unequal recombination underlies allelic copy number variation of NLRs (Kuang et al. 2004; Nagy and Bennetzen 2008; Baurens et al. 2010; Richter et al. 1995; Smith and Hulbert 2005; Q. Sun et al. 2001).

Clusters of NLR genes have been among the most difficult genomic regions to assemble alongside other repeat-rich genomic sequences including the centromeric, telomeric, and rDNA regions (Read et al. 2020). Long-read genome assemblies have finally resolved contiguous sequences across these challenging regions and are beginning to revolutionize our understanding of their function and evolution (Naish et al. 2021; Fultz et al. 2023). Characterization of the evolutionary dynamics of NLR clusters is further challenged by limitations associated with the annotation of this gene family. Many NLR gene sequences go undetected in standard gene annotation processes and thus NLR gene content is misrepresented (Bayer, Edwards, and Batley 2018). To resolve this issue, methods for sequence homology based reannotation of NLRs have been developed and have helped discover full-length NLRs that were initially overlooked (Steuernagel et al. 2020; Giuseppe Andolfo, Dohm, and Himmelbauer 2022). Complete assembly and annotation of NLR genes and gene clusters can now serve as a springboard for unraveling the mutational and evolutionary forces shaping the genomic content and organization of NLRs in plant genomes.

Long-lived species of trees sense and respond to pathogens throughout their lives, but their long generation times provide a further disadvantage in the arms-race with their fast evolving foes. Long-term pathogen pressure may cause expansion of immune receptor gene families and a broad survey has revealed expansion of RP, RK, and NLR gene families in tree species (Ngou et al. 2022). Here we explore the evolutionary dynamics of NLRs in a series of perennial tree species primarily in the genus *Citrus*. Interspecific hybridization of multiple species in this genus, as well as close relatives, has given rise to many cultivated types of citrus, including mandarins, sweet oranges, and lemons (Guohong Albert Wu et al. 2018). Like many perennial tree crop species, *Citrus* species and interspecific hybrids are not inbred, and variation in gene content between homologous chromosomes (i.e. haplotypes) may promote ectopic and unequal recombination. In *Arabidopsis thaliana*, unequal recombination during meiosis was 6 fold higher when a segment of inserted sequences was hemizygous (X.-Q. Sun et al. 2016). This, in combination with the small genome size of *Citrus* species (∼360 Mb), make it a good model for exploring the evolution of complex genomic clusters, including those with NLRs.

Here we assembled three high-quality diploid genome assemblies for a mandarin, a lemon, and a sweet orange. These three assemblies were combined with eight other citrus genome assemblies that span the genus *Citrus* and close relatives (G. Albert Wu et al. 2014; X. Wang et al. 2017; L. Wang et al. 2018; Peng et al. 2020; B. Wu et al. 2023; Nakandala et al. 2023) to characterize the patterns and processes shaping evolution of NLR gene content across 9.8 million years of evolution (Peng et al. 2020). First, the identification of NLRs was unified across all 11 genome assemblies. This library of NLR genes was then used to address two main questions. The first question focused on the genomic patterns of NLR genes in *Citrus* genomes. Despite the rapid evolution of NLR clusters, to what extent is gene content conserved in clusters across millions of years of evolution? The second question addressed the mechanistic processes driving these genomic patterns. Can non-homologous recombination explain the genomic distribution and gene content of NLR clusters? We find that despite increased nucleotide divergence at NLR gene clusters, substantial synteny between clusters could be identified. Importantly, we observed preferential expansion of phylogenetically related NLRs within clusters and conservation of gene family identity between syntenic clusters. Despite long-term conservation of gene family content between clusters, NLR clusters are in continuous genomic flux. Even so, recent lineage-specific duplications were identified that lend further support for the role of unequal recombination in shaping genomic variation in these regions.

## Results

### Haploid-resolved genome assembly for a mandarin, lemon, and sweet orange

To explore the evolutionary dynamics of nucleotide-binding leucine-rich repeat (NLR) genes across the genus we mined 15 haplotypes of citrus and citrus relatives for genes containing canonical NLR domains. The focal genomes include haploid representations of mandarin (*C. reticulata*), pummelo (*C. maxima*), and citron (*C. medica*), the three species that gave rise to cultivated citrus types through a series of interspecific hybridization events (Guohong Albert Wu et al. 2018), as well as papeda (*C. ichangensis*), Australian round lime (*C. australis*), and trifoliate orange (*Poncirus trifoliata*). With the exception of *C. ichangensis, C. reticulata,* and *C. medica* most genomes were assembled to the chromosome level, with the nine largest scaffolds representing between 78% and 91% of the predicted haploid genome size (Seker, Tuzcu, and Ollitrault 2003; Nakandala et al. 2023; Ollitrault et al. 1994).

As a complement to published citrus genomes, we produced haploid-resolved chromosome level assemblies for three cultivated citrus types - an ancient mandarin ‘Sun Chu Sha Kat’, a Lisbon lemon ‘Limoneira 8A’, and a sweet orange ‘Washington navel’ (Figure 1 A,B,C). The ‘Limoneira 8A’ and ‘Washington navel’ assemblies were also phased, with maternal and paternal haplotypes delineated based on knowledge of their ancestry. For example, lemons are derived from a hybridization event between citron and sour orange (a hybrid with mandarin and pummelo ancestry) and the citron haplotype of each chromosome was identified based on its similarity to the citron genome. Similarly, sweet oranges are interspecific hybrids between mandarin and pummelo and the two haplotypes were delineated based on alignment to each ancestral genome following the nomenclature for the ‘Valencia’ assembly (B. Wu et al. 2023). For all three assemblies, contiguous de novo assemblies were produced based on high coverage PacBio HiFi sequence reads (‘Washington navel’ = 40X per haplotype, ‘Limoneira 8A’ = 72X; and ‘Sun Chu Sha Kat’ = 82X) (Figure 1 A,B,C). The contig N50 exceeded 1 Mb for all genomes (‘Washington navel’ = 26.18 Mb, ‘Limoneira 8A’ = 27Mb; and ‘Sun Chu Sha Kat’ = 22Mb). De novo assemblies were then scaffolded with Hi-C to produce chromosome-level assemblies. The 18 largest scaffolds (9 for each haplotype) of each assembly represent 83-86% of the predicted genome size per cultivar. For ‘Washington navel’ the 18 largest scaffolds were composed of only 44 de novo contigs, highlighting the high level of contiguity of HiFi assembled citrus genomes. The statistics of the de novo and scaffolded assemblies for each cultivar are summarized in Table S1. Assembly quality and completeness were estimated using two metrics, the percentage of single-copy orthologous genes captured in the assembly (BUSCO) and k-mer based estimates of base accuracy and assembly completeness (Merqury) (Figure S1) (Rhie et al. 2020; Manni et al. 2021). More than 97% of single-copy orthologous genes contained in the BUSCO eudicot database (n=2,326) were detected across all assembled haplotypes (‘Sun Chu Sha Kat’=97.8%, 98.6%; ‘Limoneira 8A’=98.4%, 98.4%; ‘Washington navel’=98.5%, 98.4%). Similarly, Merqury quality values (QV) based on k-mer estimates of assembly completeness indicated high quality and completeness of each genome (‘Sun Chu Sha Kat’=71.0, 71.8; ‘Limoneira 8A’=60.04, 62.8; ‘Washington navel’=72.77, 71.99). The long terminal repeat (LTR) Assembly Index (LAI) (Ou, Chen, and Jiang 2018), which evaluates the contiguity of intergenic and repetitive regions of genome assemblies based on the intactness of LTR retrotransposons (LTR-RTs) of ‘Washington navel’, ‘Sun Chu Sha Kat’ and ‘Limoneira 8A’ genome assembly averaged 20.1, 22.58, 22.78 respectively, indicating a high level of contiguity of the repetitive and intergenic regions compared to other published genomes (Figure 1B, Figure S2).

**Figure 1.**
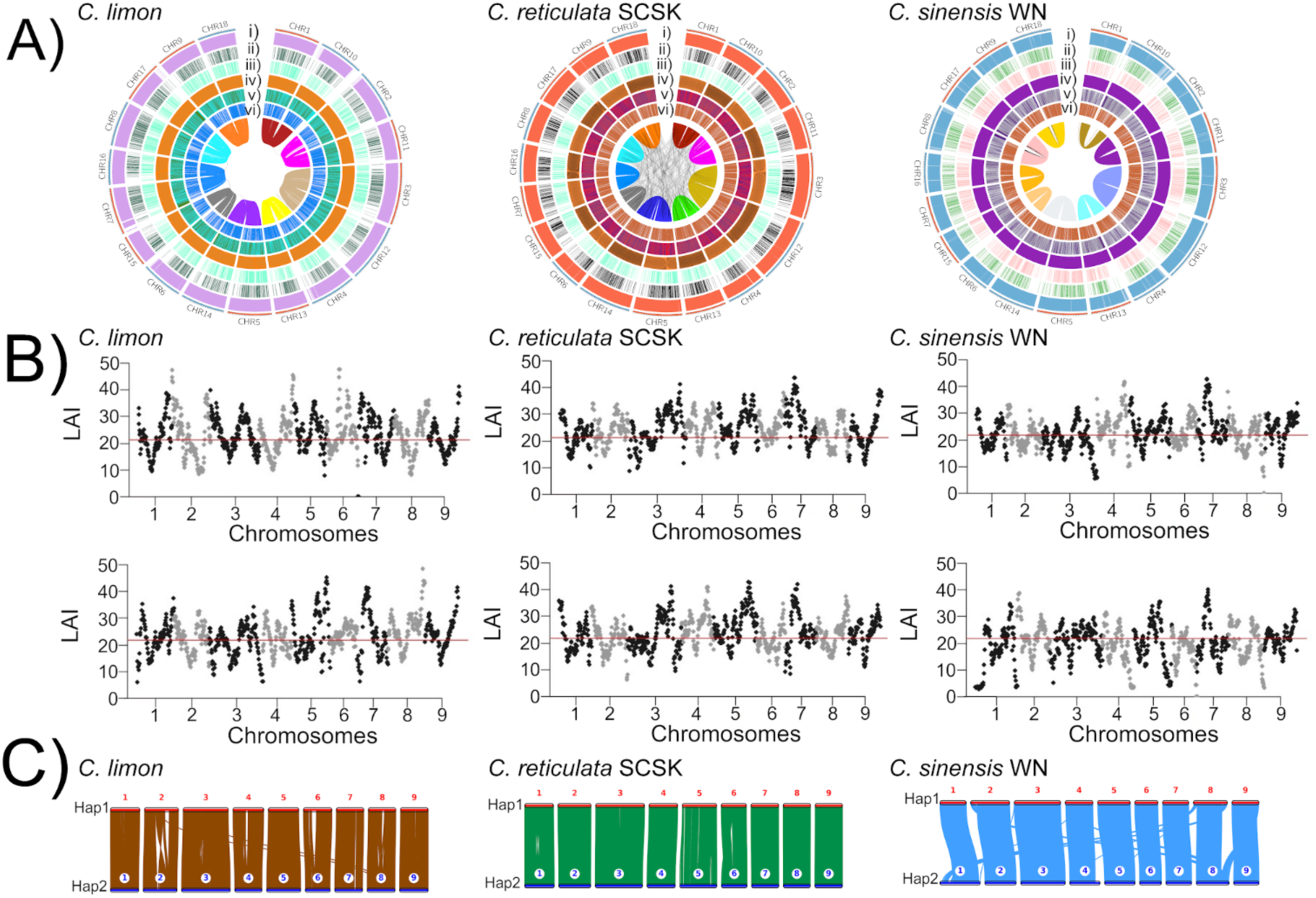
The landscape of three haplotype-resolved genome assemblies. Genomic landscape of chromosome-scale assemblies for **(A)** *C. limon* ‘Limoneira 8A’, *C. reticulata* ‘Sun Chu Sha Kat’, and *C. sinensis* ‘Washington navel’. Beginning with the outermost layer, tracks on the circos diagrams represent: (i) Gene density; (ii) Copia transposons (iii) Gypsy transposons (iv) GC percent (v) GC content; (vi) DNA transposons; (5) gene density per Mb **(B)** Evaluation of genome assembly completeness using the LTR Assembly index (LAi) for each haplotype (top, bottom). The LAI was calculated in 3 Mb sliding windows with 300 Kb steps. The average genomic LAI is plotted as a red horizontal line. **(C)** The collinearity between haplotypes in each genome is shown. Nomenclature: *C. limon* ‘Limoneira 8A’ (*C. limon*), *C. reticulata* ‘Sun Chu Sha Kat’ (*C. reticulata* SCSK), *C. sinensis* ‘Washington navel’ (*C. sinensis* WN).

Each genome was annotated using a combination of *in silico* gene prediction and empirical RNA-seq evidence. For ‘Sun Chu Sha Kat’ and ‘Limoneira 8A’ empirical evidence included PacBio long-read Iso-Seq libraries from leaf tissue, while RNA-seq data for ‘Washington navel’ was produced from leaf and fruit tissues using Illumina short-reads. Annotation completeness was also estimated by benchmarking of single-copy orthologs (BUSCO). For ‘Sun Chu Sha Kat’ a total of 51,385 and 55,798 gene models were annotated for haplotype 1 and haplotype 2, respectively (BUSCO: haplotype 1= 97.7%; haplotype 2= 97.0%; combined=97.7%). For ‘Limoneira 8A’, 44,382 and 46,298 gene models were annotated for haplotype citron and haplotype sour orange (BUSCO: haplotype sour orange= 92.1%; haplotype citron= 92.5%; combined=96.9%). A smaller number of genes were annotated in ‘Washington navel’, 31,191 genes for haplotype A (mostly mandarin ancestry) and 32,447 genes for haplotype B (mostly pummelo ancestry). Despite the reduction in genes annotated for ‘Washington navel’, the BUSCO estimates of annotated completeness were high (BUSCO: haplotype A=98.3%; haplotypeB=98.4%; combined= 98.9%).

The high level of completeness and contiguity of the three assembled genomes revealed large-scale haplotype-specific structural variation, including multi-megabase long stretches of satellite repeats. Haplotype-specific variation in satellite abundance has previously been demonstrated by *in situ* hybridization, including for sweet orange which contains both mandarin and pummelo ancestry (He et al. 2020; Song et al. 2023). Whole-genome alignment of the two ‘Washington navel’ haplotypes revealed long segments of haplotype-specific DNA segments on chromosomes 1, 2, 4, 8, and 9 (Figure S3). These regions had previously been repeat-masked by short repeat sequences (< 200 bp) and so we annotated tandem repeat sequences for these regions and across the entire genome using the software TRASH (Wlodzimierz, Hong, and Henderson 2023). The consensus sequence for 31 unique tandem repeats was identified in the ‘Washington navel genome’, with 20 tandem repeats spanning 11.26 Mb of sequence in haplotype A and 16 tandem repeats spanning 16.34 Mb in haplotype B. The majority of tandem repeats were assembled in long arrays on chromosome 1A, 1B, 2A, 4A, 4B, 8A, 8B, 9A, and 9B and these regions coincide with haplotype-specific differences observed in the alignment of the two haplotypes and haplotype-specific banding observed by *in situ* hybridization in sweet orange (Figure 1C, S3). Sequence alignment and phylogenetic analysis of the consensus sequence for 92 tandem repeats identified in ‘Washington navel’, ‘Sun Chu Sha Kat’, and ‘Limoneira 8A’ revealed that repeat sequences were typically distinct between ‘Limoneira 8A’ and the other two genomes (Figure S4). Chromosome-level assemblies for these three citrus cultivars revealed long-stretches of satellite sequence repeats that had not been clearly resolved in previously published assemblies.

### The spectrum of NLR diversity in citrus

Nucleotide-binding leucine-rich repeat (NLR) genes are rapidly evolving genes involved in the intracellular perception of pathogen effectors. Capturing the repertoire of NLR genes is challenging for at least two reasons. First, they tend to be clustered in complex genomic regions and the sequence of these regions is often not fully resolved in genome assemblies (Read et al. 2020; W. Wang et al. 2021). Second, it is well documented that NLRs are difficult to annotate, even in well assembled genomes (Jupe et al. 2013; Giuseppe Andolfo et al. 2014). This is due, in large part, to the fact that NLRs are commonly masked prior to gene annotation, preventing their inclusion in final gene annotation sets (Bayer, Edwards, and Batley 2018). To characterize the extent that these factors have impacted the discovery and annotation of NLR genes in citrus, we first used protein-homology based-prediction to identify putative NLR genes in 15 citrus haplotypes, including those from ‘Sun Chu Sha Kat’, ‘Limoneira 8A’, and ‘Washington navel’. NLRs were required to have at least two of the three canonical NLR domains, a central nucleotide-binding (NB-ARC) domain, a C-terminal leucine-rich repeat domain, and either a coiled-coil (CC) or Toll-like, Interleukin-1 receptor, Resistance protein (TIR) N-terminal signaling domain. The number of predicted NLRs identified by a protein-domain-search varied extensively between genomes, ranging from fewer than 100 to more than 500 NLRs per haplotype (Figure 2A; Table S5). This level of interspecific variation in NLR gene content is substantial, and potentially due to technical challenges in annotating NLR genes discussed above. To unify NLR discovery across new and existing genomes, we re-annotated NLR genes in all 15 haplotypes using a couple of sequence-homology based methods, HRP and NLR-annotator (Steuernagel et al. 2020; Giuseppe Andolfo, Dohm, and Himmelbauer 2022). With this approach, a library of candidate NLR sequences was developed including full-length NLRs previously identified in each genome. This library included 6,819 NLR sequences which were then used to query each genome for previously unannotated NLR sequences (Supplementary File 1). Depending on the genome, up to 5.1X additional complete NLR gene sequences were discovered (Figure 2A; Table S6; Supplementary File 2). This included 2,041 putative NLRs with all three canonical domains that were absent from the original annotations. NLR proteins lacking an N-terminal signaling domain have been shown to function in plant immunity, and so this class of predicted NLRs (NL) was included in the final set for each genome (Table S6). An example of a region in *C. australis* where we were able to recover a set of 16 NLRs missing from the original genome annotation is shown in Figure 2B. In addition to the three canonical domains in NLR proteins, which were enriched in the expected regions of the full-length NLR protein sequence (Figure 2C), a subset of NLRs may have additional integrated protein domains. We identified 266 NLRs with integrated domains across the 15 haplotypes, ranging from 7 NLRs in *P. trifoliata* up to 25 NLRs in *C. ichangensis* (Figure 2D). A total of 38 integrated domains were identified and were located primarily on the C-terminal region (25 domains), but they were also present on the N-terminal region (8 domains), or occasionally on both the N-and C-terminal regions of the protein (5 domains) (Figure 2D). ‘Importing beta binding domain’ (PF01749) was the most frequent integrated domain (40 appearances) and the gag-polypeptide of LTR copia-type (PF14223) domain also occurred frequently (26 appearances).

**Figure 2.**
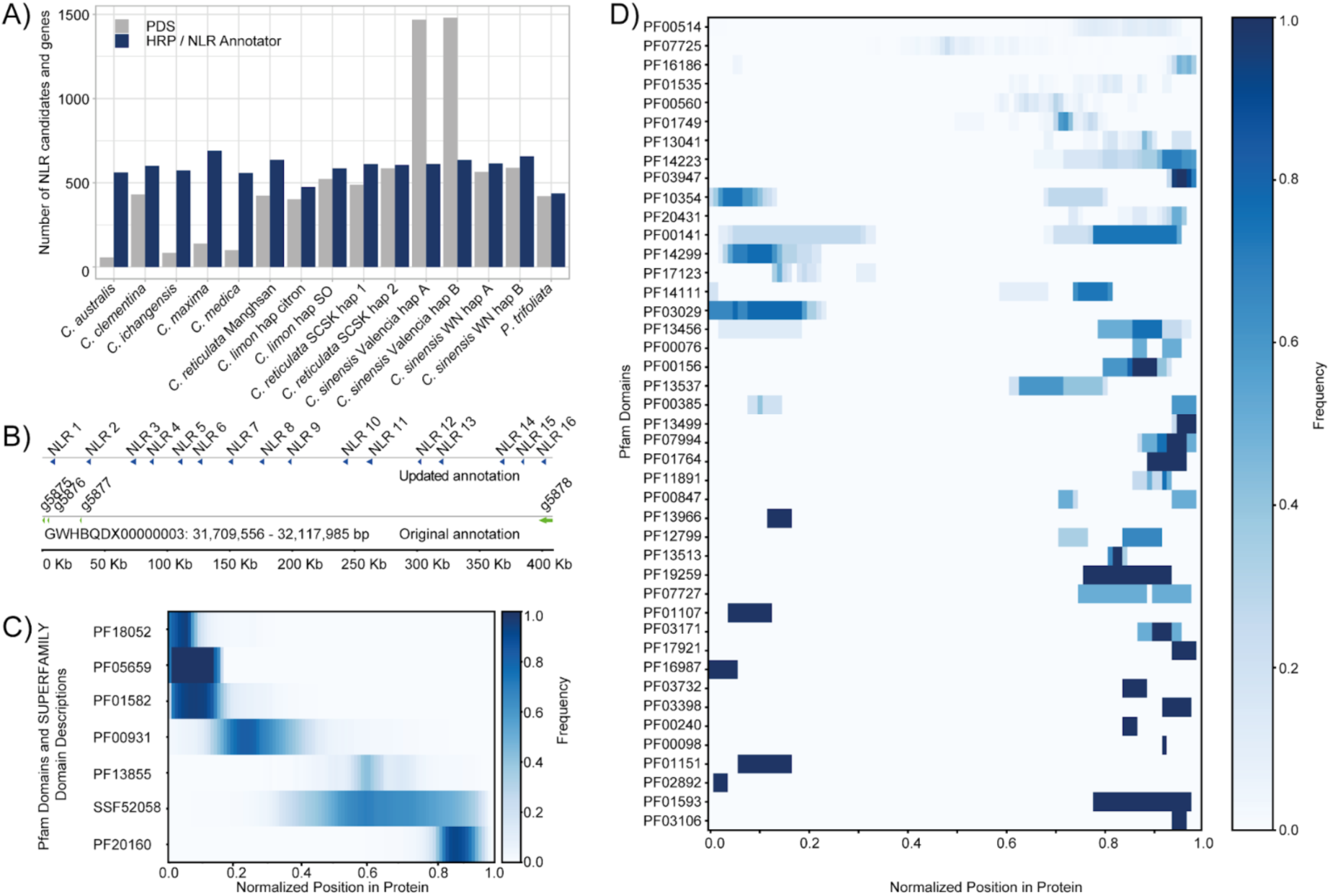
Improved annotation of NLRs in 15 citrus haplotypes. **(A)** The number of NLR identified using protein-domain searches (PDS) of published annotations (grey) and improved annotation of NLRs using Homology-based R-gene Prediction (HRP) and NLR-Annotator (blue). **(B)** Schematic representation of a cluster of 16 NLRs (blue arrows) that were missing in the published annotation from *C. australis*. Other gene categories in the published annotation are depicted as green arrows. **(C)** Normalized physical position and frequency for 6 Pfam and 1 Superfamily domains in the improved reannotation (HRP) of NLRs from all 15 haplotypes. The following domains are shown: PF18052 (CC), PF05659 (RPW8), PF01582 (TIR), PF00931 (NB-ARC), PF13855 (LRR), SSF52058 (L domain-like), and PF20160 (C-JID). **(D)** Normalized physical position and frequency for integrated domains in the improved reannotation (HRP) of NLRs from all 15 haplotypes. Nomenclature: *C. limon* ‘Limoneira 8A’ (*C. limon*), *C. reticulata* ‘Sun Chu Sha Kat’ (*C. reticulata* SCSK), *C. sinensis* ‘Washington navel’ (*C. sinensis* WN), sour orange (SO).

The focal genomes include representatives from the three ancestral citrus species as well as citrus relatives and interspecific hybrids. Citrus has evolved from an ancient paleohexaploid ancestor and has undergone chromosomal fissions and fusions to form nine chromosomes (Feng et al. 2021). ‘Washington navel’, ‘Sun Chu Sha Kat’ and ‘Limoneira 8A’ along with other *Citrus* species have undergone only one ancient whole-genome duplication (WGDs) event that is common to all eudicots (Figure S5). Unlike other eudicots, no lineage specific WGD diversification is observed in *Citrus* (Figure S5). To determine if these genomes capture NLR diversity in citrus, we first evaluated the number of new gene families discovered upon the addition of each new genome. A total of 21,050 orthogroups containing two or more genes were identified using OrthoFinder v2.5.4 (Emms and Kelly 2019). Between 91.1% and 99.9% of proteins in each genome were assigned to orthogroups (Figure 3A). A maximum-likelihood phylogeny of the 15 haplotypes was constructed using 5,254 single-copy ortholog sequences, resulting in species relationships consistent with previous studies (Figure 3B). As expected, the haplotypes of ‘Washington navel’ are most closely related to pummelo (*C. maxima*) and mandarin (*C. reticulata* and *C. clementina*). One haplotype of ‘Limoneira 8A’ is most closely related to citron (*C. medica*) and the other haplotype, which originates from sour orange, has pummelo and mandarin ancestry (Bao et al. 2023). The number of orthogroups discovered with each additional genome begins to plateau around 12 (Figure 3D), suggesting that the 15 haplotypes represent the majority of common genes in this set. A similar pattern was observed for orthogroups including NLR proteins (Figure 3G). Most orthogroups contained genes from either many or few genomes, with the largest number of multi-gene orthogroups containing genes from all 15 haplotypes (Figure 3C). Next, gene families were defined as orthogroups containing two or more genes and predicted to have at least one gene copy in the most recent common ancestor of all taxa (Mendes et al. 2020). *Poncirus trifoliata* diverged from the genus *Citrus* 9.8 million years ago (Peng et al. 2020) and using this species as an outgroup a total of 11,436 gene families were identified, including 54% of the 21,050 orthogroups. There is substantial variation in the number of gene family members per species (Figure 3E), with 214-502 gene families significantly expanded or contracted per genome (Table S7).

**Figure 3.**
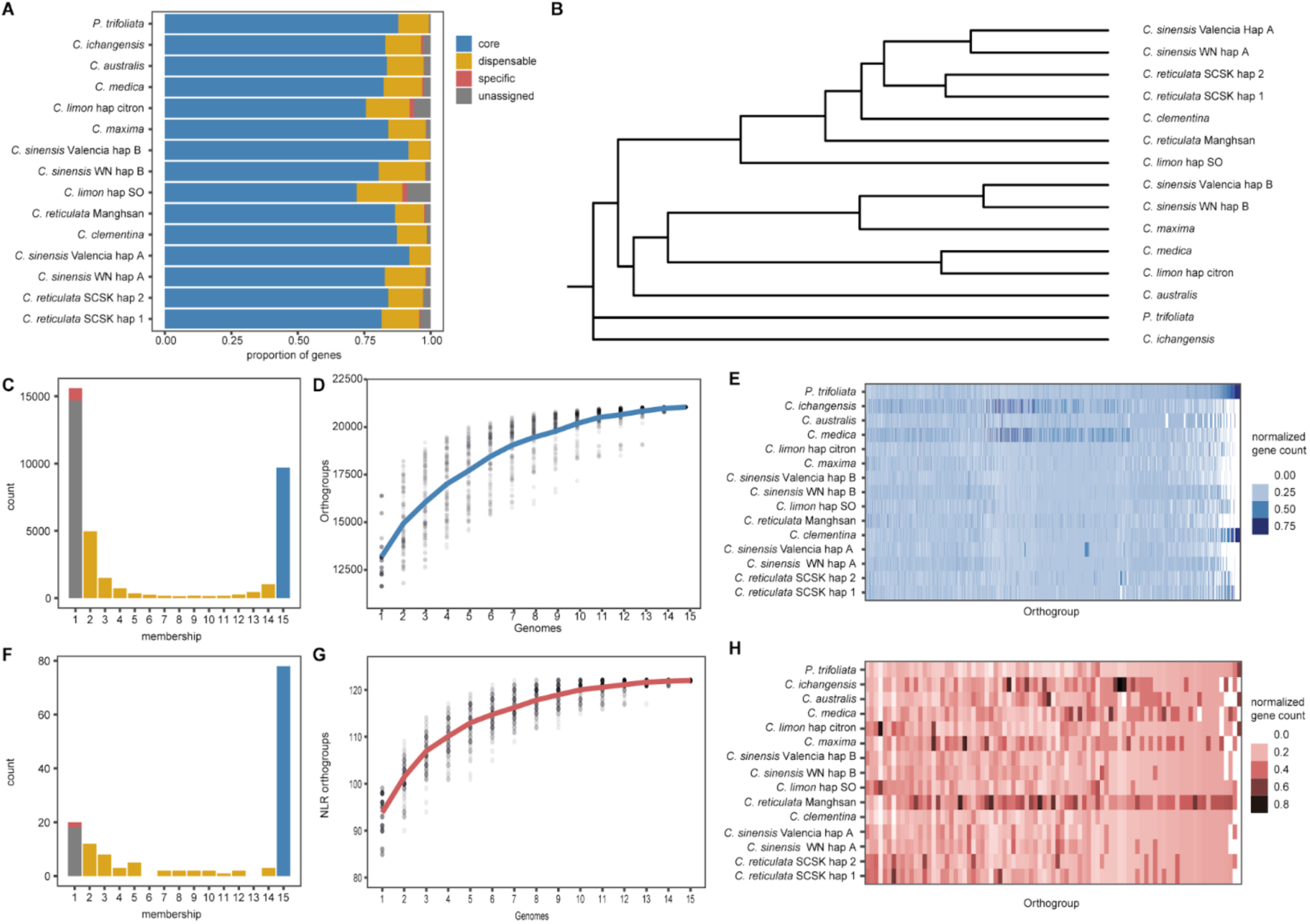
Identification of gene families across 15 citrus haplotypes. **(A)** The percentage of genes in each genome assigned to orthogroups. Core genes are those with orthologs in all 15 haplotypes, dispensable genes were identified in fewer than 15 haplotypes, and specific genes were identified in a single haplotype **(B)** Maximum-likelihood tree constructed using single copy ortholog protein sequences. **(C)** The number of orthogroups containing members from one or more haplotypes. There may be one or more members from each haplotype. **(D)** A rarefaction curve of the number of orthogroups discovered with an increasing number of haplotypes. The solid line represents the mean of 100 permutations. **(E)** A heatmap of scaled gene count per gene family. Gene counts are scaled within gene families. **(F)** The number of NLR-containing orthogroups containing members from one or more haplotype. There may be one or more members from each haplotype. **(G)** A rarefaction curve of the number of NLR-containing orthogroups discovered with an increasing number of haplotypes. The solid line represents the mean of 100 permutations. **(H)** A heatmap of scaled gene count per NLR-containing gene family. Gene counts are scaled within gene families. Nomenclature: *C. limon* ‘Limoneira 8A’ (*C. limon*), *C. reticulata* ‘Sun Chu Sha Kat’ (*C. reticulata* SCSK), *C. sinensis* ‘Washington navel’ (*C. sinensis* WN), sour orange (SO).

Next we focused on the 122 orthogroups (0.6% of orthogroups) and 73 gene families (0.6% of gene families) that contained one or more predicted NLR genes. The number of NLR orthogroups discovered also increased with the addition of each haplotype and approached saturation around 12 haplotypes, similar to the plateau observed for all orthogroups (Figure 3G). This indicates that the majority of common NLRs are represented in this set, but that inclusion of additional genomes will facilitate the discovery of additional NLRs that occur at lower frequencies across the genus. Approximately 98% of NLR orthogroups consisted of members from 2 or more haplotypes (Figure 3F), with 78 orthogroups containing members from all haplotypes. These orthogroups contained 84% of the HRP-annotated NLRs, including 82% of the full-length CNLs, 86% of the full-length TNLs, and 87% of the full-length RNLs. Similar to all genes, the number of members of each NLR orthogroup varied across species (Figure 3H), with only a few NLR orthogroups significantly expanded or contracted per species (Table S8).

NLRs exhibit high levels of protein sequence variation, including those evolving under both directional and balancing selection (Van de Weyer et al. 2019). Previous work in Arabidopsis, Brachipodium, and maize identified clades of NLRs that show elevated amino acid sequence diversity in each species (Prigozhin and Krasileva 2021; Prigozhin et al. 2024). Using a similar methodology, we characterized the number and type of highly variable NLRs (hvNLR) across the genus *Citrus*. A total of 11122 NLRs, including the 15 focal haplotypes, were phylogenetically grouped into 350 refined clades (Figure 4). Shannon entropy was used as a measure of NLR amino acid diversity and hvNLR clades were defined as those with at least 10 amino acid residues exceeding an entropy score of 2. We identified 41 hvNLR clades (12% of the NLR clades) containing 49% of the analyzed NLRs, ranging from 192 to 359 hvNLR per haplotype (Table S9). The hvNLRs included 65% of the full-length CNLs, 46% of the full-length TNLs, and no RNLs. This result is consistent with previous studies demonstrating that hvNLRs are distributed throughout CNL and TNL clades in *Arabidopsis thaliana* (Prigozhin and Krasileva 2021).

**Figure 4.**
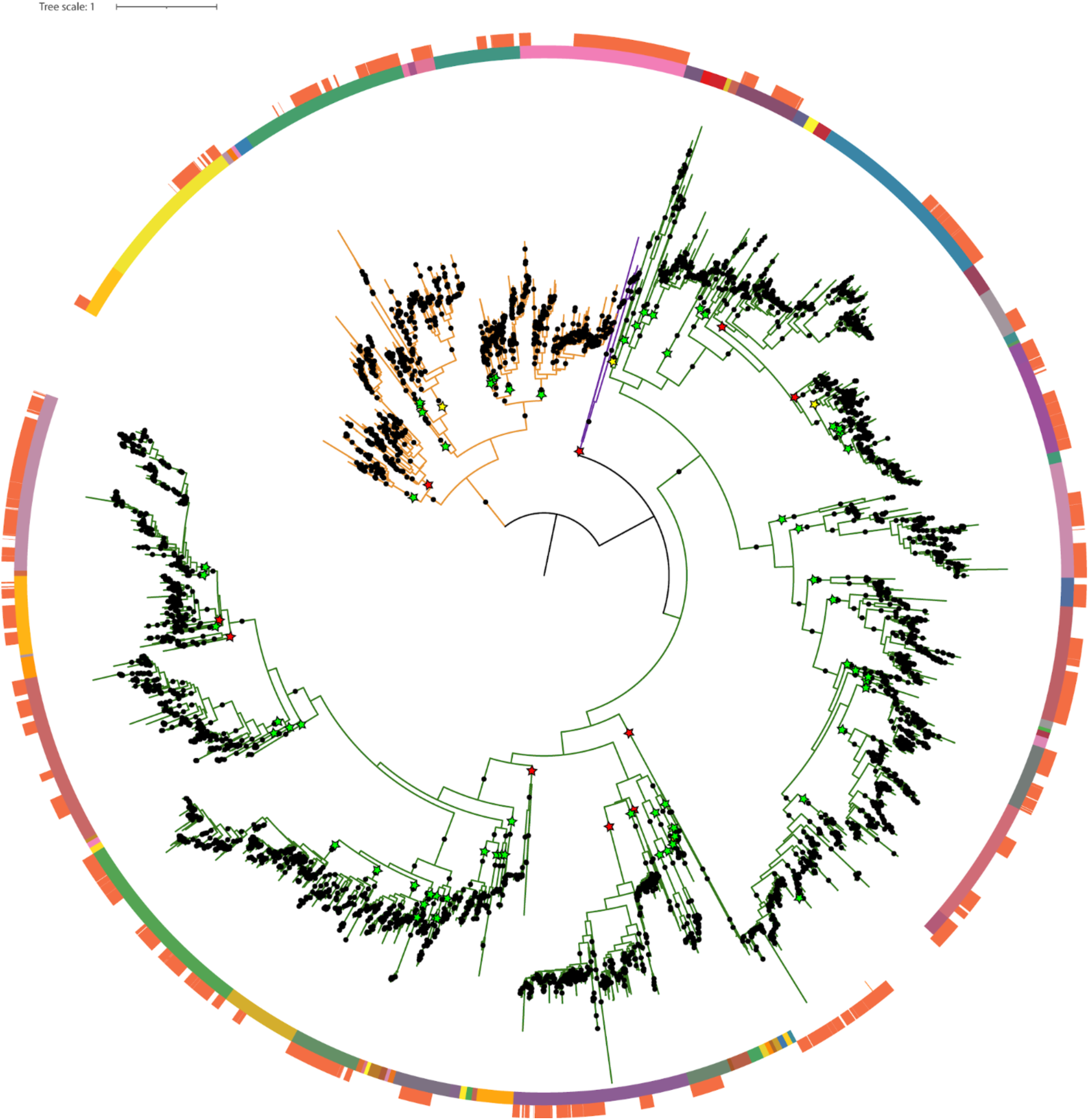
Phylogeny of citrus NLRs showing RNL, TNL, and CNL subfamilies as purple, orange, and green branches respectively. The tree was split on branches highlighted with stars (green for bootstrap support > 90, yellow > 70, and red otherwise); the resulting initial clades are shown as colored ranges on the inside circle. Initial clades were refined and hvNLRs defined as sequences with refined clade protein alignments with 10 or more residues at 2 bits or higher entropy. The hvNLRs are highlighted in orange on the outer circle. Bootstrap values of 70 and higher are shown as dots. Tree scale is average substitutions per site.

### The genomic clustering of NLRs

The genomic distribution of NLR gene sequences is non-random, with NLRs often residing in complex genomic clusters (Shao et al. 2014; Zhou et al. 2004; R. R. Q. Lee and Chae 2020; Woudstra et al. 2024; Q. Li, Jiang, and Shao 2021). This arrangement is hypothesized to facilitate the rapid evolution of NLR genes through expansion and contraction of clustered genes via ectopic recombination, although direct evidence of this process is limited. To understand the evolutionary dynamics of NLR gene clusters we first characterized the genomic distribution of NLRs. NLRs are distributed throughout the 18 chromosomes and are densely clustered in some genomes (Figure S6). Genome assembly breakpoints were not enriched for NLRs (Table S10). Across all 15 haplotypes, the median intergenic distance between NLRs was 16,245 bp (Figure S7). NLR intergenic distance did vary substantially between scaffolds (In *C. limon* ‘Limoneira 8A’ haplotype citron, min = 13,781 bp; max = 1,193,208 bp), highlighting their non-random genomic distribution. For the 12 chromosome-level assemblies, 76.9% of NLRs (n = 5,370) resided on three of the nine chromosomes and these three chromosomes represent less than 40% of assembled genome length in any of the 12 haplotypes.

With this information we then systematically identified NLR clusters in each genome to explore the degree of conservation of cluster sequence and gene membership over nearly 10 million years of evolution. Previous studies have delineated NLR clusters using a wide range of inter-NLR distances (Jupe et al. 2012; Ameline-Torregrosa et al. 2008; R. R. Q. Lee and Chae 2020). The relationship between number of clusters and inter-NLR distance was used to determine the optimal distance for cluster determination in citrus (Figure 5A, S8). Based on this, adjacent NLRs in each of the 15 haplotypes were grouped into physical clusters if the maximum inter-NLR distance was 50 kb or less. With this criteria, between 80 and 127 clusters of two or more NLR genes were identified per genome (Table S11). The outgroup species *P. trifoliata* had the lowest number of NLR clusters (n=80). The genome with the largest number of NLR clusters (*C. reticulata Mangshan* (n=127) was not scaffolded to the chromosome level, which may result in fragmentation of larger clusters. In all of the chromosome-level genome assemblies, 77% or more of NLRs are contained in a cluster (Table S12). Similar to previous studies in *Arabidopsis thaliana* and *Hordeum vulgare*, most clusters (51.9%) contain three or fewer NLRs (Jiao and Schneeberger 2020; Q. Li, Jiang, and Shao 2021). Clusters with large numbers of NLRs (>10) were also identified, including 9.98% of clusters in the surveyed genomes (Figure 5B). In some cases, clusters with as many as 35 NLRs were identified (Figure 5B). Although most NLRs in a single cluster belong to the same classification (CNL, TNL, RNL, or NL), around one-third of clusters (ranging from 20% in *C.limon* ‘Limoneira 8A’ haplotype citron to 38% in *C. australis*) contain NLRs across classification types (CNL, TNL, RNL, or NL). Most clusters with multiple classes of NLRs contained one or more members from a single type (CNL, TNL, or RNL) together with an NL. Only eleven clusters (0.75% of all clusters) contained a combination of full-length NLRs classes (i.e. CNL - RNL or CNL - TNL) (Table S13).

**Figure 5.**
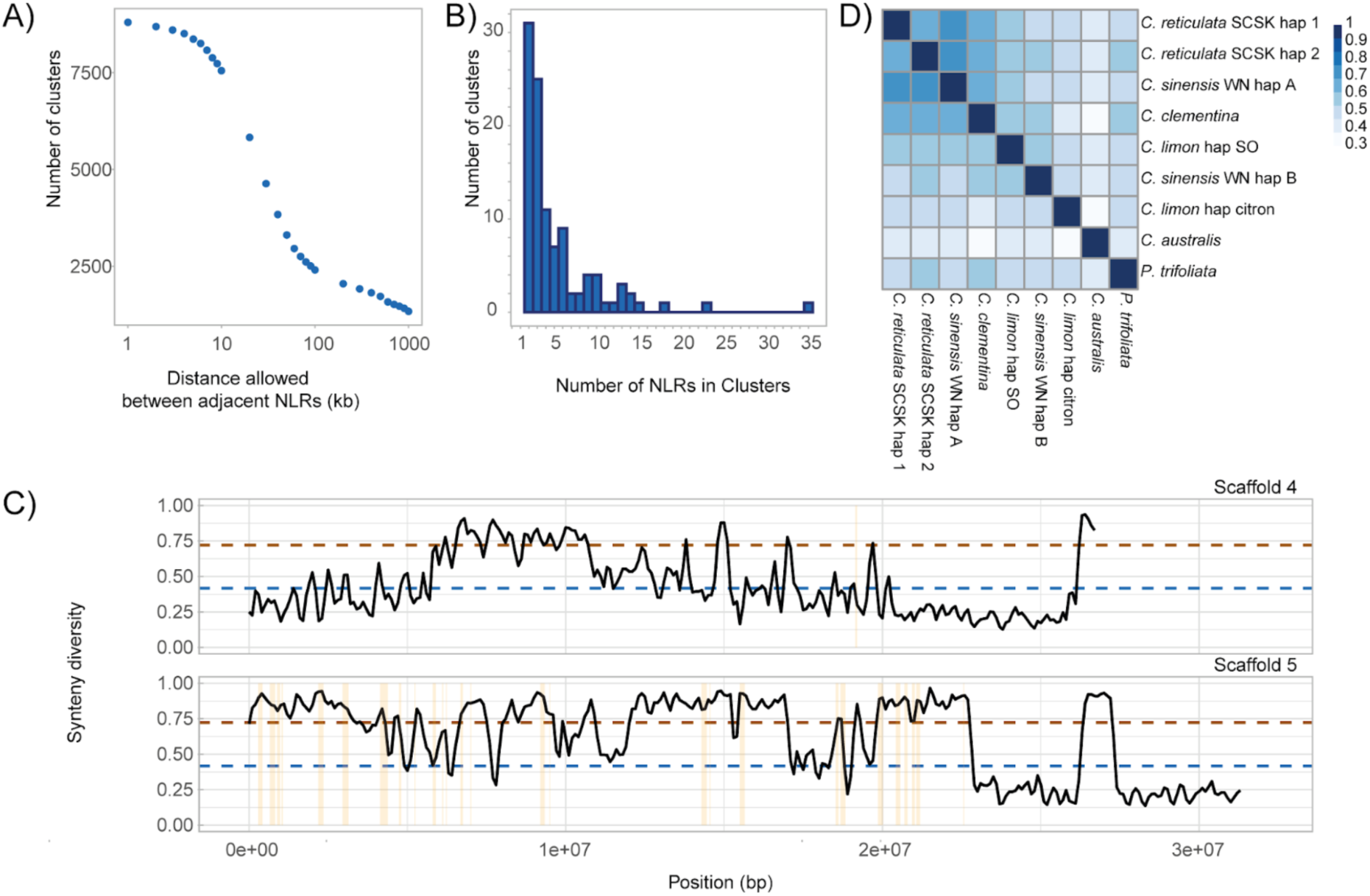
NLRs are clustered and clusters are syntenic between genomes. **(A)** The relationship between physical distance between adjacent NLRs and the number of identified clusters. NLR clusters were determined based on a 50 Kb intergenic distance. **(B)** The distribution of the number of NLRs per cluster. **(C)** Synteny diversity scores for two representative scaffolds. *P. trifoliata* was used as the reference genome. Scaffold 4 is an example of a scaffold with few NLR clusters. In contrast, many NLR clusters reside on Scaffold 5. Yellow bars highlight the physical location of NLR clusters in *P. trifoliata*. Synteny diversity scores were calculated for 200 Kb windows with a 100 Kb step. The dotted lines represent synteny diversity scores of 0.417 (blue) and 0.722 (brown). **(D)** A heatmap of Jaccard scores for pairwise synteny of NLR clusters across nine chromosome-level genome assemblies. Nomenclature: *C. limon* ‘Limoneira 8A’ (*C. limon*), *C. reticulata* ‘Sun Chu Sha Kat’ (*C. reticulata* SCSK), *C. sinensis* ‘Washington navel’ (*C. sinensis* WN), sour orange (SO).

With a unified annotation of NLR genes and subsequent NLR cluster identification, we were able to explore the evolutionary dynamics of genomic regions underlying NLR clusters. First we evaluated the degree of sequence collinearity across all chromosome-level genome assemblies. These assemblies include haplotypes derived from mandarin (*C. reticulata* ‘Sun Chu Sha Kat’ haplotypes 1 and 2, *C. sinensis* ‘Washington navel’ haplotype A, *C. clementina*), pummelo (*C. sinensis* ‘Washington navel’ haplotype B, *C. limon* ‘Limoneira 8A’ haplotype sour orange), and citron (*C. limon* ‘Limoneira 8A’ haplotype citron), as well as for citrus relatives including *C. australis* and *P. trifoliata*. Based on pairwise whole-genome alignments, a metric to describe the average degree of collinearity or synteny (synteny diversity or *π_syn_*) in genomic windows was estimated based on (Jiao and Schneeberger 2020). Using pairwise alignments for nine genomes/haplotypes, we estimated *π_syn_* in 200 Kb windows (100 Kb step) and visualized genomic patterns of *π_syn_* relative to *P. trifoliata* (Figure 5C, S9). Regions syntenic across all nine genomes are reflected by a *π_syn_* score of 0. There is a wide range of *π_syn_* scores across all nine citrus chromosomes (Figure 5C). This metric was developed to describe the average degree of synteny in a series of *A. thaliana* genomes (Jiao and Schneeberger 2020) and compared to these intraspecific genomic comparisons, there are several regions in our interspecific analysis of citrus genomes that contain large rearrangements or other structural variants in at least three out of nine of the genomes analyzed (score >= 0.583). To confirm that NLR genes are more likely to reside in complex genomic regions with increased *π_syn_*, we determined *π_syn_* in windows overlapping the 80 clusters of NLR genes in *P. trifoliata* (Figure S10). Indeed, the average *π_syn_*for these 80 clusters was significantly larger than randomly sampled genomic regions of similar size (*p* < 0.0001, based on 10,000 permutations).

Although NLR clusters are located in genomic regions with reduced synteny, we find that pairwise synteny at NLR clusters is still substantial. Syntenic pairs of NLR clusters were defined as clusters with at least 15% of aligned nucleotide sequences along their genomic lengths. In total, 1,967 pairs of syntenic NLR clusters were identified from whole-genome alignments of the nine chromosome-level assemblies (Supplementary File 3). Average pairwise synteny at NLR clusters was high (Figure 5D; Table S14, mean Jaccard score = 0.51) and haplotypes with shared ancestry (i.e. mandarin or pummelo) tended to have larger proportions of syntenic NLR clusters. For example, the mostly mandarin derived *C. sinensis* WN haplotype A and mandarin *C. reticulata* SCSK haplotype 2 were syntenic at a majority of NLR clusters (Jaccard score = 0.72). Lower synteny was observed in more distant comparisons, including *C. australis* and *C. limon* ‘Limoneira 8A’ haplotype citron (Jaccard score = 0.29). There is also variation in NLR gene content and cluster synteny within haplotypes in a single accession. ‘Sun Chu Sha Kat’ is a mandarin with no evidence of interspecific admixture, and its two haplotypes, *C. reticulata* SCSK haplotype 1 and 2, are not syntenic at all clusters of NLRs (Jaccard score = 0.68). Despite sequence divergence at genomic clusters enriched with NLR genes (Figure 5D), there is considerable conservation of NLR cluster identity indicating that NLRs tend to retain their genomic location.

### Dynamics of gene content at syntenic clusters of NLRs

Pairwise conservation of NLR cluster identity was used to investigate the dynamics of NLR gene cluster evolution. There is no consensus regarding the mechanism underlying rapid evolution of NLR clusters and there is at least some evidence for ectopic recombination (interlocus), unequal recombination (intralocus), and transposon-mediated gene capture (David et al. 2009; Jupe et al. 2012; D. Leister et al. 1998) in generating NLR clusters with variable gene content, including the creation of chimeric NLRs with novel functions (Nagy and Bennetzen 2008; Q. Sun et al. 2001; Smith and Hulbert 2005; Baurens et al. 2010; Richter et al. 1995; Kuang et al. 2004). NLRs are often clustered as tandem duplicates (Jupe et al. 2012; Guo et al. 2011; G. Andolfo et al. 2013), and sequence duplications are prone to unequal recombination (John G. Jelesko et al. 2004; Wicker, Yahiaoui, and Keller 2007) leading to gain or loss of intervening sequence (Nagy and Bennetzen 2008; Cossu et al. 2017; Wicker, Yahiaoui, and Keller 2007). Local expansion and contraction of NLR sequences would result in clusters of paralogous NLRs and careful examination of specific NLR clusters has revealed that clustered NLRs share an evolutionary history (Baurens et al. 2010; Smith and Hulbert 2005). We had previously characterized orthologous NLR gene families (Figure 3) and used this analysis to determine if NLR clusters tended to contain paralogs derived from the same orthogroup. Indeed, a majority (72%) of NLR clusters across the 15 surveyed haplotypes contained genes from a single orthogroup (Figure 6A). In fact, 92.3% of NLR clusters contain only 1 or 2 NLR orthogroups. Multi-orthogroup clusters of NLR tended to have more genes and be longer than single-orthogroup clusters (Gene number: W = 192100, p= 4.58^-14^; Cluster length: W = 201976, p=7.94^-10^) (Figure S11). Additionally, in multi-orthogroup clusters, a single orthogroup tended to be overrepresented in the cluster, suggesting preferential local amplification of a select orthogroup (Figure 6B). Among 439 multi-orthogroup clusters, overrepresentation of a single orthogroup was detected for 70 NLR clusters (Chi-squared goodness of fit, p<=0.05) (Figure 6B). Multi-orthogroup clusters with preferential enrichment of one orthogroup included more genes (median = 11 genes) compared to those where no preference was detected (median = 4 genes). Finally, NLRs located at syntenic clusters (Figure 5) are enriched for members of the same ortholog groups, suggesting that cluster gene content is conserved in addition to genomic location (Figure 6C) (1,646 out of the 1,967 (83.6%) syntenic pairs of NLR clusters). Together this suggests that local duplication of NLR gene sequences is the major driver of gene content evolution at NLR clusters, with preferential amplification of single orthogroups in larger gene clusters and conservation of gene content at syntenic genomic regions.

**Figure 6.**
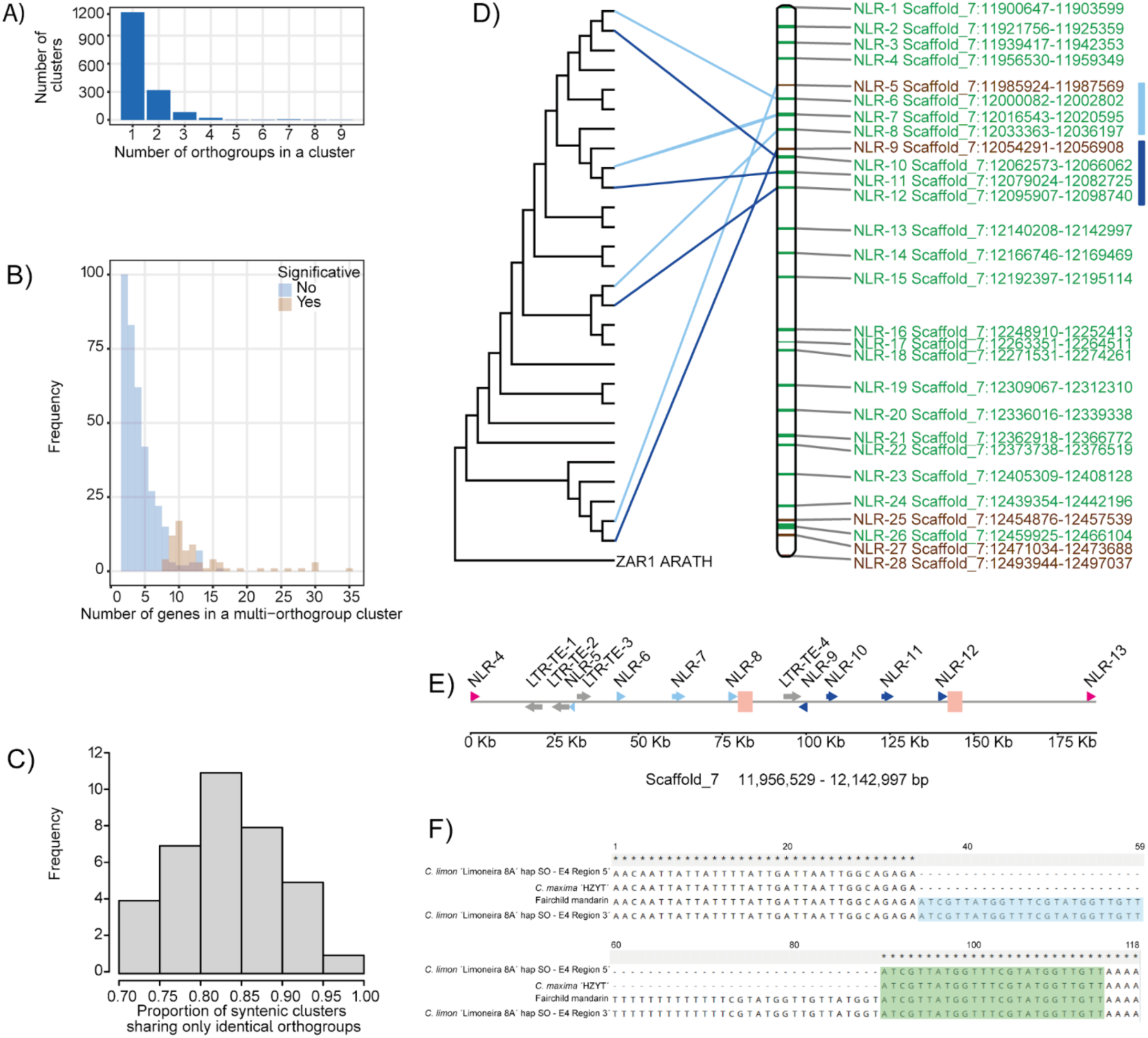
Gene content variation in NLR clusters results from expansion and contraction of single orthogroups. **(A)** The number of NLR-containing orthogroups per NLR cluster. **(B)** The distribution of the number of NLRs per multi-orthogroup NLR cluster. Clusters without over-representation of a single orthogroup (blue) tend to have fewer NLRs than clusters with significant over-representation of a single orthogroup (brown). **(C)** The distribution of the proportion of syntenic NLRs clusters with identical orthogroup membership. Pairwise comparison of syntenic clusters performed for the 9 chromosome-level haplotypes. **(D)** A phylogenetic tree of NLRs in clusters 80 and 81 in *C. limon* ‘Limoneira 8A’ haplotype sour orange (left) and their physical position on scaffold 7 (right), rooted with the HOPZ-ACTIVATED RESISTANCE 1 (ZAR1) NLR from *Arabidopsis thaliana*. The IDs of NLRs from the most frequent orthogroup are labeled in green. The IDs of NLRs from additional orthogroups are labeled in brown. NLRs that reside in the recent duplication identified in *C. limon* ‘Limoneira 8A’ haplotype sour orange are highlighted by the light and dark blue lines connecting the NLR position in the phylogenetic tree with the physical location in the cluster for a set of duplicated NLRs. **(E)** A schematic representation of the duplicated region in cluster 80 in *C. limon* ‘Limoneira 8A’ haplotype sour orange. Duplicated NLRs are depicted in light and dark blue arrows and non-duplicated NLRs as pink arrows. LTR-TE are depicted as gray arrows. The “E4” regions are denoted by the pink boxes. (F) Nucleotide alignment from a section of the “E4” region showing evidence of unequal recombination. The alignment includes the sequences of the 5’ and 3’ E4 regions from *C. limon* ‘Limoneira 8A’ haplotype sour orange as well as sequence to represent pummelo (C. maxima ‘Huazhouyou-tomentosa’ (HZYT)) and mandarin (‘Fairchild’) ancestry. The 25-bp region (light blue) is also found after the 55-bp indel (light green).

Next we searched for sequence signatures at NLR clusters that might elucidate the mechanism of local duplications associated with gene expansion. Sequence duplications can arise from transposon-mediated capture of gene sequences and insertion of these sequences in new genomic locations. In plants, gene capture by both class I and class II transposons can occur (Muyle et al. 2021). Proliferation of long terminal repeat (LTR) retrotransposons moves through an RNA intermediate, and gene sequences captured by LTRs will lack introns (Cerbin and Jiang 2018). We noted several examples of LTR TEs neighboring NLRs (Figure S12). To systematically evaluate the contribution of LTR-mediated duplication of NLRs, 842 intronless NLRs across 12 chromosome-level assemblies were identified (not including, *C. ichangensis*, *C. reticulata*, *C. medica*). An intron-containing donor NLR gene was identified for only 125 of the intronless NLRs (Table S15). The majority of these pairs were found between genomes or haplotypes, suggesting that either the donor or duplicate gene copy was lost. There were 14 examples of intron-containing donor and intronless duplicate NLRs in the same haplotype (Table S15) which may be examples of more recent retroduplication. Although there are examples where NLR duplication was associated with retroduplication, these examples are not sufficient to explain massive regional expansion of clustered NLR genes.

Repair of double stranded breaks can lead to unequal recombination when a nearby, but non-allelic, repair template is used, or ectopic recombination when the repair template is from a distant genomic location. In both cases, non-allelic repair of double stranded breaks will result in sequence expansion of one chromatid or homolog, depending on which is used to provide the template filler DNA sequence (Wicker, Yahiaoui, and Keller 2007), and sequence loss on the corresponding chromatid/homolog. We reasoned that recent duplications within NLR clusters were most useful for resolving sequences at the duplication breakpoints, and thus useful for identifying sequences associated with non-homologous recombination. To identify recent duplications, self-alignments were performed for the NLR clusters found in haplotypes from ‘Washington navel’ (*C. sinensis* ‘Washington navel’ haplotype A, *C. sinensis* ‘Washington navel’ haplotype B), ‘Limoneira 8A’ (*C. limon* ‘Limoneira 8A’ haplotype citron; *C. limon* ‘Limoneira 8A’ haplotype sour orange), or ‘Sun Chu Sha Kat’ (*C. reticulata* ‘Sun Chu Sha Kat’ haplotype 1; *C. reticulata* ‘Sun Chu Sha Kat’ haplotype 2). Self-alignments revealed a spectrum of sequence similarity at NLR clusters. A majority of clusters (52.3% of 615 alignments) included secondary alignments that spanned at least 10% of their genomic length (Figure S13). Secondary alignments that made up 50% or more of the region were also identified (11.5% of 615 alignments), which is a pattern consistent with more recent sequence duplications (Figure S14). In a few rare cases, no secondary alignments were identified, even for a cluster with 12 NLRs.

Based on this analysis, recent duplications were identified for two haplotypes of a syntenic NLR cluster, *C. sinensis* ‘Washington navel’ haplotype B and *C. limon* ‘Limoneira 8A’ haplotype sour orange. There is substantial variation in NLR gene content at this cluster across the surveyed genomes, ranging from 11 NLRs in *C. australis* to 35 in *C. sinensis* ‘Washington navel’ haplotype B (Figure S15). Recent partial duplication of this cluster in *C. sinensis* ‘Washington navel’ haplotype B and *C. limon* ‘Limoneira 8A’ haplotype sour orange were supported by phylogenetic analysis of NLR protein sequences (Figure S16). In both cases, multiple neighboring NLRs were tandemly duplicated (Figure 6D and E, Figure S17). In *C. sinensis* ‘Washington navel’ haplotype B, this cluster (C89-WN_B) contains 35 NLRs and a single orthogroup has preferentially expanded (Figure S15). There are two recent duplications in *C. sinensis* ‘Washington navel’ haplotype B. The first duplication includes a120.7 Kb 5’ region and a 117.4 Kb 3’ region, with 155.3Kb of intervening sequencing. There is 99.3% sequence identity between the 5’ and 3’ regions of this duplication which includes a total of ten NLRs (five in each region) and five intact LTR RTs (three in the 5’ region and two in the 3’ region). The second duplication in *C. sinensis* ‘Washington navel’ haplotype B includes eight NLRs, four in a 106.7 Kb 5’ region and four in a 99.8Kb 3’ region with 99.7% sequence identity between the regions. There is 302 Kb of intervening sequence. There is also a 51 Kb overlap between the 5’ regions of the first and second duplications in *C. sinensis* ‘Washington navel’ haplotype B and this region contains two NLRs (Figure S15). Due to the complexity of duplications in this haplotype, we were unable to pinpoint sequence signatures that are associated with unequal or ectopic recombination. While repetitive sequences are found throughout the duplicated regions and their flanking sequences, there were no repetitive sequences specifically associated with the duplication break points.

The duplication in *C. limon* ‘Limoneira 8A’ haplotype sour orange spanned eight NLRs, with four NLRs each in a 60.5 Kb 5’ region and a 54.6 Kb 3’ region. These duplicated regions were 98.9% identical and were separated by 7.8 Kb of intervening sequence. The larger size of the 5’ region is due to the insertion of an intact Ty-1-copia LTR retrotransposon approximately 508 thousand years ago, based on divergence of the LTR sequences (SanMiguel et al. 1998). The *C. limon* ‘Limoneira 8A’ haplotype sour orange contains ancestry from both mandarin and pummelo and sequences derived from both of these ancestors were found in the duplicated region. The 5’ duplicated region (which includes NLRs 5-8) and some sequences upstream of this region are 99.6% identical to a mandarin haplotype in the ‘Fairchild’ genome assembly (Diaz et al. 2024). Importantly, the similarity with ‘Fairchild’ haplotype extends past NLR 5 to a 4.3 Kb sequence called “E4” (Figure S18). We could not identify a haplotype with exact sequence identity to the entire 3’ region, but multiple pummelo haplotypes (Huazhouyou-tomentosa, Majia, and Majiayou) contain sequences that are more than 98% identical to the sequence from a portion of “E4” in the 5’ region up to the stop codon of NLR 9 in the 3’ region (Figure S18) (Zheng et al. 2023; Huang et al. 2023; Lu et al. 2022). Although a haplotype with high sequence similarity across the entire 3’ region of this duplication could not be identified, we suspect that assembly of additional haplotypes from pummelo will resolve this issue.

Unequal recombination is initiated by the formation of a double stranded break. Double stranded breaks can result from DNA damage (Pfeiffer, Goedecke, and Obe 2000) or transposon integration (Pfeiffer, Goedecke, and Obe 2000), and there are some regions of the genome that are prone to double stranded breaks for reasons that are not clear (Ji et al. 2022). The breakpoints of ectopic recombination events in monocot genomes included sequences associated with both TEs and other repetitive sequences (Wicker, Buchmann, and Keller 2010). Because the “E4” flanked both the 5’ and 3’ regions and was also located upstream of sequences identified in pummelo genomes with high similarity to the 3’ region, we searched this region for signatures of unequal recombination. Indeed, sequence alignment of “E4” from the 5’ and 3’ regions with “E4” from ‘Fairchild’ and ‘Huazhouyou-tomentosa’ revealed a 55 bp indel consistent with unequal recombination, with a 25 bp sequence repeated on the 5’ internal border and 3’ region flanking the deletion (Figure 6F, Figure S19) (Wicker, Yahiaoui, and Keller 2007). We propose that unequal recombination between a mandarin haplotype similar to ‘Fairchild’ and a pummelo haplotype at the “E4” regions of these two haplotypes led to the duplication of four NLR genes in *C. limon* ‘Limoneira 8A’ haplotype sour orange. The “E4” region includes 4,357 bp of sequence with no annotated genes. However, sequence analysis revealed that positions 1,870-2,898 have high protein sequence identity (84%) to the LRR domain of some NLRs in citrus, suggesting that a portion of “E4” encodes a truncated NLR. The 55 bp indel is 1,434 bp upstream of the truncated NLR. Although expansion and contraction of NLRs in clusters is a common feature of plant genomes, it is extremely challenging to pinpoint the exact mechanism promoting gene content variation in these regions. By focusing on recent sequence duplications in a highly variable, syntenic cluster of NLRs we were able to identify at least one case resulting from unequal recombination between two divergent haplotypes.

## Discussion

Poor assembly of complex genomic regions and incomplete annotation of NLR genes has limited investigations into the evolutionary dynamics and genomic shuffling of NLR gene clusters. Here, we have integrated high-quality, contiguous genome assemblies with unified NLR annotations to characterize the genomic patterns of NLRs across the genus *Citrus* and explore the mechanistic processes shaping NLR genomic content.

### The patterns: NLRs arranged in genomic clusters with shared evolutionary history

We first defined the NLR gene content for 11 citrus accessions, including four diploid assemblies where both haplotypes were resolved. Prior to reannotation there was substantial variation in the number of NLRs per genome. Unification of NLR annotations using homology-based prediction reduced this variation, with similar numbers of NLRs detected per genome in the TNL, CNL, RNL, and NL categories. These 15 haplotypes also represent the majority of NLR gene families in *Citrus*, with saturation in NLR gene content observed in rarefaction analysis (Figure 3G). This comprehensive set of NLR genes identified across the three foundational species of cultivated citrus (mandarin, pummelo, and citron), their interspecific hybrids, and relatives enabled the exploration of the evolution of this gene family over nearly 10 million years of evolution.

We find the majority of NLRs in each lineage reside in genomic clusters composed of paralogous genes, typically from a single gene family, suggesting that genomic clustering of NLRs arose through local duplication events. Even large NLR clusters are limited to genes derived from one or a few gene families. In NLR clusters with multiple gene families there is preferential enrichment of a single gene family lineage, with significant enrichment evident when clusters contain 8 or more NLRs (Figure 6B). The fact that NLR genes are clustered in genomes has been known since the first NLRs were cloned (Whitham et al. 1994; Bent et al. 1994; Mindrinos et al. 1994). Early identification of NLR clusters also revealed that neighboring NLRs tend to share an evolutionary history. Here we show that the evolution of NLR clusters across a genus is associated with preferential expansion and contraction of specific gene families, even in clusters that contain multiple gene families (Figure 6B). This required identification of NLR clusters with synteny between genomes. Although NLR clusters reside in regions of the genome with elevated divergence, synteny between clusters could still be identified (Figure 5D). In any pairwise comparison, synteny could be determined for an average of 70% of NLR clusters per genome. There are 18 clusters where synteny could be inferred for all lineages and the patterns of NLR expansion and contraction determined across the genus. These 18 clusters revealed the dynamic nature of NLR cluster evolution. In one case, the number of NLRs in a syntenic cluster varied from 19 in the outgroup *P. trifoliata* to 35 in ‘Washington navel’ orange, an interspecific hybrid containing both mandarin (*C. reticulata*) and pummelo (*C. maxima*) ancestry. Surprisingly, the lineage with the smallest number of NLRs was *C. australis* (11 NLRs) followed by *C. limon* ‘Limoneira 8A’ haplotype citron (16 NLRs) and stepwise gains and losses of NLR number across the phylogeny were not clear. Similar trends were observed for other clusters where synteny could be determined across the genus. This suggests that lineage specific evolution of NLR gene content in syntenic clusters is rapid. Previous work in *Arabidopsis thaliana* also suggested that intraspecific copy number variation of NLR genes was restricted to certain members of multi-gene clusters (R. R. Q. Lee and Chae 2020) and groups of orthologous NLR tend to reside in just one genomic region (Teasdale et al. 2024). This analysis was based on sequence capture of NLRs across a panel of *A. thaliana* accessions and subsequent alignment to a single reference, making it difficult to determine if NLR copy number variation reflects local or global variation in NLR cluster gene content. We extend this previous work by clearly demonstrating that NLR copy number variation reflects local genomic variation in NLR cluster content, a finding that favors mechanisms that promote local expansion and contraction of genomic sequences in NLR clusters.

### The processes: Evidence for unequal recombination promoting expansion and contraction of NLRs in genomic clusters

Next we sought to address the mechanistic processes that promote local clustering of paralogous NLRs. To do so, we focused on instances of recent NLR duplication, reasoning that the genomic signatures of the processes driving duplication were more likely to be evident at or surrounding recent duplicates. Gene duplication can result both from transposon-mediated gene capture and proliferation and mechanisms of non-homologous recombination (Cerbin and Jiang 2018). It was unlikely that transposon-mediated proliferation would be the major driver of local cluster variation because this would require transposons with captured NLR sequences to integrate repeatedly in the same local region. Even so, we looked for signatures of transposon-mediated capture and duplication of NLRs. Based on phylogenetic analysis, very few recently duplicated NLRs were identified within a genome/haplotype where one copy lacked intron sequences, a signature of retrotransposition (Cerbin and Jiang 2018). Overall, regional clustering and patterns of local duplication of NLRs are unlikely to be driven primarily by retrotransposition in citrus.

Non-homologous recombination is favored as a mechanism driving local sequence duplication, including at NLR gene clusters (Ronald 1998). During non-homologous recombination multiple possible non-allelic templates can be used to repair a double-stranded break. Ectopic recombination refers to examples where the repair template is located at a distant genomic location. Ectopic recombination is the process associated with gene movement in genomes (Wicker, Buchmann, and Keller 2010) and there is at least one example where ectopic recombination led to a novel cluster of NLR genes (David et al. 2009). Detailed analysis of genes in non-collinear genomic locations in *Brachypodium distachyon* and *Oryza sativa* revealed signatures associated with gene duplication, including ectopic recombination (Wicker, Buchmann, and Keller 2010). Sequences flanking duplicates including both transposon-associated sequences and other types of repeats (Wicker, Buchmann, and Keller 2010). Based on this, it was suggested that doubled-stranded breaks, which often occur in repeats and transposons (Pfeiffer, Goedecke, and Obe 2000), were the trigger for ectopic recombination, not the flanking sequences itself (Wicker, Buchmann, and Keller 2010). A transgenic assay for ectopic recombination in tobacco provided direct evidence that double stranded breaks in TEs, and not the TE sequence itself, were required for ectopic recombination (Shalev and Levy 1997).

Unequal recombination also uses non-allelic templates to repair double stranded breaks, but, unlike ectopic recombination, those templates are located in closer proximity to the double stranded break. Examples of unequal recombination generating novel alleles at loci with tandem sequence duplications are abundant, including at NLR clusters (Nagy and Bennetzen 2008; Q. Sun et al. 2001; Smith and Hulbert 2005; Baurens et al. 2010; Richter et al. 1995; Kuang et al. 2004). We identified recent sequence duplications at the NLR clusters identified in citrus, with a focus on duplications in the contiguous assemblies of ‘Washington’ navel, ‘Limoneira 8A’, and ‘Sun Chu Sha Kat’. Detailed investigation of a large syntenic NLR cluster revealed recent duplications in two haplotypes, two in *C. sinensis* ‘Washington navel’ haplotype B and another in *C. limon* ‘Limoneira 8A’ haplotype sour orange. In both cases, multiple neighboring NLRs were duplicated along with flanking sequences. The breakpoints surrounding the duplication in *C. limon* ‘Limoneira 8A’ haplotype sour orange have sequence signatures associated with unequal recombination, including a 15 bp repeat flanking the border of a deletion. Similar signatures could not be identified for the duplications in *C. sinensis* ‘Washington navel’ haplotype B. In most syntenic NLR clusters sequence identity outside of coding regions was limited and it was not possible to reconstruct the sequence of duplication and deletions at syntenic NLR clusters. Even so, unequal recombination is the most likely process to generate the genomic patterns of clustered NLR genes within and between genomes.

Factors that influence unequal and ectopic recombination have been investigated in both natural and transgenic systems. Ectopic recombination is higher for hemizygous inserted sequences in transgenic assays (X. Sun et al. 2008; X.-Q. Sun et al. 2016) and also for hemizygous natural alleles of the Cf locus in tomato, a cluster of receptor proteins associated with resistance to the fungal pathogen *Cladosporium fulvan (Parniske et al. 1997; Wulff et al. 2004; Takken et al. 1998)*.

There are also examples of hemizygosity reducing rates of unequal recombination (Yandeau-Nelson et al. 2006). Large portions of the genomes of heterozygous perennial crops are hemizygous, and this holds true for the three diploid assemblies presented here. The haplotypes of NLR clusters that are syntenic between the A and B haplotypes of ‘Washington’ navel are also dissimilar, with variation in both sequence and NLR content. Double-stranded breaks in hemizygous sequences of NLR clusters may promote unequal recombination and support the local expansion and contraction of paralogous NLR genes. There are also sequences that are prone to unequal recombination. Sequence homology is expected to promote unequal recombination, but some studies indicate that unequal recombination takes place across a spectrum of sequence similarities (Nagy and Bennetzen 2008). Numerous ectopic recombination events map to arrays of leucine-rich repeats (LRR) (Nagy and Bennetzen 2008; Parniske et al. 1997; Michelmore and Meyers 1998). These regions are also prone to non-homologous end joining (NHEJ) where ends of DNA sequence fragments are joined based on very short regions of sequence homology (Raina et al. 2021) and which can expand or contract LRR arrays (Nagy and Bennetzen 2008). The 3’ breakpoints of both the 5’ and 3’ regions that make up the recent duplication in *C. limon* ‘Limoneira 8A’ haplotype sour orange includes sequence with homology to an LRR domain, although no other flanking N-terminal or NB-ARC domains were detected. A signature of non-homologous recombination was evident in the alignments between these two regions, with a 15 bp sequence flanking a deletion containing the same 15 bp sequence on the interior border of the deletion (Wicker, Yahiaoui, and Keller 2007). This deletion was ∼1 Kb downstream of the truncated LRR domain. Although it is easy to invoke unequal recombination at NLR clusters to explain patterns of clustered genes with shared evolutionary histories, unequal recombination was initially expected to lead to sequence homogenization, where paralogs within a cluster have higher sequence identity to one another than orthologs (Michelmore and Meyers 1998; Ronald 1998). There is still work needed to resolve this issue, but there is at least some evidence that unequal recombination can generate allelic series, leading to diversification and not homogenization of duplicated sequences (John G. Jelesko et al. 2004).

## Conclusion

Plant genomes are in a constant state of change. They tolerate repeated whole genome duplication events, large-scale genomic rearrangements, and genomic conflict associated with replication and proliferation of transposable elements. Clusters of NLR genes are prime examples of this genomic flux. These regions have elevated levels of sequence divergence and reduced collinearity between genomes (Jiao and Schneeberger 2020). More rapid sequence evolution and genomic shuffling in NLR clusters may help long-lived plants, which lack adaptive immune systems, to keep pace with pathogen evolution, which typically have much shorter generation times. Through careful reannotation of NLR genes in citrus we demonstrate that NLR clusters preferentially contain paralogous genes and identify recent duplications which support a role for ectopic recombination in generating genome-wide patterns of regional clustering of genes with a shared evolutionary history.

## Methods

### Genome sequencing

Mature leaf tissue was collected from a single *C. sinensis* ‘Washington Navel’ orange, *C. limon* ‘Limoneira 8A’, and *C. reticulata* ‘Sun Chu Sha Kat’ accessions at the University of California, Riverside Givaudan Citrus Variety Collection in Riverside, CA. For ‘Washington Navel’, high molecular weight DNA was isolated using the protocol “Preparing Arabidopsis Genomic DNA for Size-Selected ∼20 kb SMRTbell^TM^ Libraries” with the addition of a sorbitol wash step following nuclei isolation.

For ‘Limoneira 8A’ and ‘Sun Chu Sha Kat’, nuclei were isolated from plant tissues and HMW DNA was extracted from the nuclei using the Circulomics Nanobind Plant Nuclei Big DNA kit (Pacific Biosciences, Menlo Park, CA, USA). For all accessions, libraries were constructed and sequenced using the Pacific Biosciences Sequel II platform by the DNA Technologies and Expression Analysis Core at the UC Davis Genome Center, supported by NIH Shared Instrumentation Grant 1S10OD010786-01. The CCS software (https://github.com/PacificBiosciences/ccs) was used to produce high-precision HiFi reads with quality above Q20. SMRTbell adapter contamination in the HiFi reads was checked using cutadapt (v2.10) (Martin 2011).

#### Scaffolding with Hi-C for ‘Washington Navel’ HiC

A Dovetail HiC library was prepared in a similar manner as described previously (Lieberman-Aiden et al. 2009). Briefly, for each library, chromatin was fixed in place with formaldehyde in the nucleus and then extracted Fixed chromatin was digested with DpnII, the 5’ overhangs filled in with biotinylated nucleotides, and then free blunt ends were ligated. After ligation, crosslinks were reversed and the DNA purified from protein. Purified DNA was treated to remove biotin that was not internal to ligated fragments. The DNA was then sheared to ∼350 bp mean fragment size and sequencing libraries were generated using NEBNext Ultra enzymes and Illumina-compatible adapters.Biotin-containing fragments were isolated using streptavidin beads before PCR enrichment of each library. The libraries were sequenced on an Illumina HiSeq X to produce 178 million 2x150 bp paired end reads.

#### Scaffolding with Hi-C for ‘Limoneira 8A’ and ‘Sun Chu Sha Kat’

Crosslinking was performed on ground tissue and nuclei were isolated following (Colt 2021). The Arima Hi-C library was constructed using an Arima-HiC Kit (Arima Genomics, CA, USA) according to the manufacturer’s instructions for Plant Tissues (A160135 v01). Libraries were generated using the Roche KAPA Hyper Prep and sequencing was done in Illumina HiSeq.

### RNA sequencing

#### Plant material, RNA extraction and IsoSeq sequencing

For ‘Washington navel’, albedo, flavedo, juice vesicles, and segment membrane from three different fruits collected from the original Parent Washington navel tree were stored at either 4°C for 24 hours or at room temperature. Similarly, three replicates were collected from young leaves, mature leaves, floral buds, and mature flowers from the original ‘Parent Washington navel’ tree. The tissue was immediately frozen with liquid nitrogen and stored until its use. RNA was extracted using Trizol according to the manufacturer’s instructions. RNA was sent to Amaryllis Nucleics (Oakland, CA) for library preparation and Illumina sequencing to generate 851.8 million 2 x150 paired-end reads.

For *C. limon* ‘Limoneira 8A’ and *C. reticulata* ‘Sun Chu Sha Kat’, total RNA was extracted from leaf tissue collected from a single tree accession at the University of California, Riverside Givaudan Citrus Variety Collection in Riverside, CA, with the RNeasy plant mini kit (QIAGEN – including on-column DNase I treatment) according to the manufacturer’s instructions. NEBNext Single cell/low input cDNA synthesis & amplification module and PacBio Iso-Seq express oligo kit were used to perform first-strand cDNA synthesis and PCR amplification. Barcoded forward and reverse primers (barcoded NEBNext Single Cell cDNA PCR Primer and barcoded Iso-Seq express cDNA PCR Primer) were used during PCR amplification of cDNA samples. Size selection was done with the BluePippin DNA Size-Selection System (Sage Science). The size-selected cDNA were used to prepare the cDNA SMRTbell libraries. DNA damage repair and SMRTbell ligation was performed with SMRTbell Express Template Prep Kit 2.0 (Pacific Biosciences, Menlo Park, CA, USA) following the manufacturer’s protocol. Finally, the SMRTbell library was sequenced on a PacBio Sequel II system (DNA Technology Core Facility, University of California, Davis, CA, USA).

The raw long-read fastq sequences were processed using the Iso-seq3 (v3.8.0) pipeline (https://github.com/PacificBiosciences/IsoSeq). Briefly, the PacBio IsoSeq raw subreads were demultiplexed using Lima (https://lima.how/). Circular consensus reads (ccs) were generated from subreads using PacBio Ccs program (https://ccs.how/). PolyA tails and concatemers were trimmed from ccs reads using Isoseq3 refine. Redundant reads were clustered based on pairwise alignment using Isoseq3 (https://github.com/PacificBiosciences/IsoSeq). Clustered transcripts were aligned to the reference genome using minimap2 (H. Li 2021). Redundant isoforms were removed using cDNA-Cupcake pipeline (http://github.com/Magdoll/cDNA_Cupcake). After filtering away isoforms, further quality control, isoform annotations and classification were done using SQANTI3 (Pardo-Palacios et al. 2024)

### Genome assembly

The genome assemblies were performed with hifiasm v0.16.0 (Cheng et al. 2022) using the Hi-C integration assembly model based on HiFi and paired-end Hi-C reads (Cheng et al. 2022). The quality and completeness of the assemblies, consisting of two sets of haplotype-resolved contigs, were evaluated by Merqury v1.3 (Rhie et al. 2020) and BUSCO v5.0.0 (Manni et al. 2021) using the eudicot database version 10. The LAI—a method to evaluate genome assembly completeness based on the quality of the assembly of repeat sequences was also run for all the genomes above using the LTR_retriever pipeline (Ou and Jiang 2018). For ‘Washington Navel’ and ‘Limoneira 8A’, the diploid assemblies were scaffolded following the 3D-DNA pipeline (Dudchenko et al. 2017). Briefly, the HiC reads were mapped against the genome assembly by BWA-MEM v0.7.17-r1188 (H. Li 2013) using Juicer v1.6 (Durand et al. 2016). Then the run-asm-pipeline.sh script from the 3D-pipeline was used to perform the scaffolding. Based on the HiC contact map, the round 0 of the scaffolding was manually corrected to fix misplaced regions using the Juicebox Assembly Tool (JBAT) (Durand et al. 2016). For ‘Washington navel’, scaffolds were assigned into two different haplotypes based on their similarity against the two haplotypes from *C. sinensis* cv Valencia (B. Wu et al. 2023). For ‘Limoneira 8A’, scaffolds were assigned into two different haplotypes based on their similarity against the *C. medica* genome assembly (X. Wang et al. 2017). For ‘Sun Chu Sha Kat’, the assembly was scaffolded into pseudo chromosomes using RagTag version 2.1.0 19 (Alonge et al. 2022).

### Genome annotation

#### Genome annotation for ‘Washington Navel’

A library of transposable elements for the two haplotypes of ‘Washington navel’ was generated by EDTA v2.0.1 (Ou et al. 2019) with the sensitive option activated. A filtering step was performed on the TE library to remove sequences encoding an open reading frame (ORF) with significant similarity against a list of 13 PFAM domains associated with NLR genes (Van Ghelder et al. 2019). Finally, the genome was soft-masked by RepeatMasker v4.1.1 (Smit, Hubley, and Green 2013-2015) using the filtered library. Transposable element related sequences (DNA transposons, long-terminal-repeat retrotransposons (LTR-RTs; Copia and Gypsy), and non-LTR retrotransposons) comprise ∼57% of the ‘Washington navel genome’ and LTR-RTs are the most abundant TEs (Table S4).

Genome annotation was performed over the soft-masked genome using the Braker V2.1.6 pipeline (Brůna et al. 2021) with hints from RNA-Seq and protein evidence. Briefly, RNA-Seq reads were generated from 12 different tissues and three replicates per tissue. Reads went through a quality filter to remove low-quality bases, polyG, short reads, and adapter sequences by fastp (S. Chen et al. 2018). Surviving reads were aligned against the Washington navel genome by HISAT (Kim et al. 2019), and the resulting bam file was used for Braker to generate intron hints.

On the other hand, hints from protein evidence were generated by prothint (Brůna, Lomsadze, and Borodovsky 2020) using a database of plant protein sequences composed of protein sequences from Uniprot/Swissprot (UniProt Consortium 2019) (accessed on 01/25/2023), OrthoDB v11 (Kuznetsov et al. 2023) (accessed on 25/11/2022), *Poncirus trifoliata (Peng et al. 2020)*, *Citrus limon* L. Burm f. (Guardo et al. 2021)*, Citrus sinensis* cv valencia (B. Wu et al. 2023), as well as the *Citrus reticulata* SCSK mandarin and *Citrus limon* L generated in this work. FInally, Braker ETP was used to predict genes with AUGUSTUS (Stanke et al. 2008) using the hints previously generated. The predicted proteins were annotated by InterProScan v5.60-92.0 (P. Jones et al. 2014). (Guardo et al. 2021)

#### Genome annotation for ‘Limoneira 8A’ and ‘Sun Chu Sha Kat’

We generated a repeat library using a combination of structure-based and homology-based approaches. Miniature inverted repeat transposable elements were analyzed using MITE-Hunter (Durand et al. 2016; Han and Wessler 2010). Long terminal repeat retrotransposons were collected using LTR-harvest and LTRdigest (Ellinghaus 2007; Steinbiss et al. 2009), followed by a filtering to exclude false positives. To collect other repetitive sequences, the genomic sequence was then masked using the long terminal repeat and miniature inverted repeat transposable element sequences. The unmasked sequence was extracted and processed by RepeatModeler (https://weatherby.genetics.utah.edu/MAKER/wiki/index.php/Repeat_Library_Construction-Advanced). The ‘Extensive de novo TE Annotator’ (EDTA) pipeline was also used to identify transposable elements (TEs) and other repetitive sequences with default parameters (Ou et al. 2019). Transposable element related sequences (DNA transposons, long-terminal-repeat retrotransposons (LTR-RTs; Copia and Gypsy), and non-LTR retrotransposons) comprise ∼45% and ∼52% of the total genome assembly of ‘Sun Chu Sha Kat’ and ‘Limoneira 8A, respectively, and LTR-RTs are the most abundant TEs (Table S2, S3).

We performed gene annotation using the MAKER3 pipeline (v.3.01.03) (Campbell et al. 2014). This included an initial round of evidence alignment, followed by two rounds of training and prediction using ab initio predictors SNAP, GeneMark-EST and Augustus (Korf 2004; Stanke et al. 2006). In the first round we provided the custom repeat library as masking evidence, taxon-specific ESTs as EST evidence, and non-redundant protein sequence database as protein evidence (421,628 proteins) downloaded from NCBI. The options est2genome=1 and protein2genome=1 were used in MAKER round one to output evidence-derived gene models for training SNAP v2013-11-29, AUGUSTUS v3.2.2 and GeneMark-ES v4.48 (Korf 2004; Stanke et al. 2006; Ter-Hovhannisyan et al. 2008). Evidence-derived gene models encoding proteins with AED scores below 0.5 were used to train SNAP, AUGUSTUS and GeneMark-ES before re-running MAKER3 to predict genes. Once the ab initio tools were trained, a new round of MAKER3 was conducted.

Putative functions of predicted proteins were identified with a BLAST v.2.2.31+ (Camacho et al. 2009) search with an E-value threshold of ≤10^−5^. Results were filtered to include only matches with >70% sequence similarity and matches longer than 50 amino acid residues. The protein motifs and domains were searched using PfamScan and InterProScan v5 based on default protein databases including ProDom, PRINTS, PfamA, SMART, TIGRFAM, PrositeProfiles, PrositePatterns, SITE, SignalP, TMHMM, Panther, Phobius, Coils and CDD (Mistry, Bateman, and Finn 2007; P. Jones et al. 2014; Attwood et al. 2012; Bru et al. 2005; Sigrist et al. 2013; Letunic, Doerks, and Bork 2015; de Lima Morais et al. 2011).

#### Non-coding gene prediction for ‘Limoneira 8A’ and ‘Sun Chu Sha Kat’

tRNAscan-SE software was used to predict the tRNAs with eukaryotic parameters. miRNAs, rRNAs, and snRNAs were detected using Infernal cmscan to search the Rfam database. The rRNAs and the corresponding subunits were annotated with RNAmmer v1.2 (Lowe and Eddy 1997; Nawrocki and Eddy 2013; Griffiths-Jones et al. 2003; Lagesen et al. 2007).

### Satellite DNA analysis

Tandem repeat identification was performed using TRASH (Wlodzimierz, Hong, and Henderson 2023) with default parameters. A neighbor-joining tree using the JC-C model with one thousand bootstrap was built by MEGA 11 (Tamura, Stecher, and Kumar 2021) for the consensus primary sequences aligned by MUSCLE (Edgar 2004). All positions with less than 90% site coverage were eliminated from the analysis.

### Synteny analysis

The BLASTP program was used to identify orthologous and paralogous genes. MCscanX79 was used to recognize syntenic blocks with parameters E_VALUE = 1e−05, MAX GAPS = 25, and MATCH_SIZE = 5. Syntenic blocks were visualized with MCscan, and chromosome lengths were not scaled (Y. Wang et al. 2012). Dot plot alignments between genome assemblies were generated and visualized using D-GENIES v1.2.0 (Cabanettes and Klopp 2018). For circos, we performed a BLAST search using gene sequences with DIAMOND87 v.2.0.9 (Buchfink, Xie, and Huson 2015). The Circos plot was generated using the Circos software (Krzywinski et al. 2009). Distributions of synonymous substitution (Ks) were constructed for each species using the primary transcripts with wgd v2 (H. Chen, Zwaenepoel, and Van de Peer 2024).

### Identification of NBS-encoding genes using annotated gene sets

We applied hmmsearch of the HMMER (v3.3) program based on the HMM corresponding to the Pfam NBS (NB-ARC) domain (PF00931) to screen and identify the NBS-encoding genes. To determine whether the candidate NBS-encoding genes encoded TIR, RPW8, or LRR motifs, the Pfam database (http://pfam.xfam.org/), SMART protein motif analyses (http://smart.embl-heidelberg.de/), and Multiple Expectation Maximization for Motif Elicitation (MEME) were used. The COILS program was also conducted to detect potential CC motifs in the NBS-encoding genes, with a threshold value of 0.9.

### De novo identification of NLR genes

Two different tools were used to annotate NLR genes. On one hand, NLR Annotator (Steuernagel et al. 2020) was run using default parameters on the citrus genomes. On the other hand, a modified approach of the HRP method (Giuseppe Andolfo, Dohm, and Himmelbauer 2022), which identifies NLR genes based on a protein domain search approach (PDS) combined with a genBlastG search (She et al. 2011), was followed. Candidate sequences from the genome annotations were combined to create a library of NLR-like proteins. Since some proteins seemed to be fused NLR-gene models (for instance, multiple N terminal, NB-ARC, LRR domains), we removed from the library sequences with more than one NB-ARC domain. Additionally, protein candidates were removed from the library if they contained either transposon or retrotransposon related domains. The filtered library was used as input to continue on the step 6 of the HRP method. Gene models were annotated by InterProScan v5.60-92.0. Gene models with no domains associated with NLR genes were removed from the analysis. Finally, gene models predicted by both NLR-Annotator and HRP, as well as full-length NLRs predicted exclusively by the HRP pipeline were kept.

### NLR clustering

NLRs were clustered based on the physical distance between adjacent NLRs in each genome, allowing up to 50 kb between adjacent NLRs to be considered part of the same cluster. Cluster nomenclature is based entirely on their order of appearance, being cluster 1 the first cluster from scaffold 1 in each assembly, and the last one is the last cluster from scaffold 9 in its respective assembly.

### NLRs with Integrated Domains

NLR with Integrated domains were defined as those who contained at least one Pfam domain that was not traditionally associated with NLRs, which includes NB-ARC, TIR, LRR, RPW8, PPR repeat, Atypical Arm repeat, Armadillo/beta-catenin-like repeat, C-JID and Rx N-terminal domain.

### Gene Family Identification

Protein sequences were first filtered to include only the longest isoform per locus. Putative NLRs identified in the genome annotation using a protein domain search approach were replaced with NLRs identified using the HRP method. Protein sequences were then clustered into orthogroups with OrthoFinder v2.5.4 (Emms and Kelly 2019) using a clustering parameter of 1. Gene families were defined as orthogroups containing two or more genes and predicted to have at least one gene copy in the most recent common ancestor to all taxa. FastTree v2.1.11 (Price, Dehal, and Arkin 2010) was used to infer an approximately-maximum-likelihood phylogenetic tree from concatenated protein sequence alignments of 5254 single-copy orthologs per genome. The resulting phylogenetic tree was then made ultrametric using r8s v1.81. Gene families and the ultrametric species tree were used as input for CAFE v5 (Mendes et al. 2020) to infer significantly expanded and contracted gene families.

### Identification of hvNLRs

hvNLRs are defined using the method described in (Prigozhin and Krasileva 2021). Shannon entropy was used as a measure of NLR amino acid diversity. Shannon entropy is calculated using the formula:

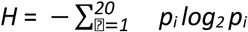

where p_i_ is the fraction of 1 of the 20 amino acids in a column of a protein sequence alignment. Highly variable NLRs (hvNLRs) were defined as NLRs with 10 or more positions exceeding 2 bit cutoff. Four additional genomes were included in the analysis of hvNLR. This included two haplotypes from the *C. limon* ‘femminello’ (Guardo et al. 2021), as well as two earlier versions of the *C. sinensis* ‘Valencia’ genome assembly (Xu et al. 2013; L. Wang et al. 2021). These four genomes were removed from further analyses because the genome assembly quality and contiguity was not sufficient.

### Determination of whole-genome synteny and pisyn

In order to identify and quantify syntenic regions across multiple chromosome-level genome assemblies, we used the synteny diversity parameter proposed by (Jiao and Schneeberger 2020). Briefly, whole genome alignments for all the possible pairwise comparisons for the genome assemblies of *C. sinensis* ‘Washington navel’ haplotypes A and B, *C. limon* ‘Limoneira 8A’ haplotypes citron and sour orange, *C. reticulata* ‘Sun Chu Sha Kat’ haplotypes 1 and 2, *P. trifoliata*, *C. australis*, and *C. clementina* were generated using nucmer v4.0.0rc1 (Marçais et al. 2018). Then, Syri v1.6.3 (Goel et al. 2019) was used to identify syntenic regions that were shared among the nine genome assemblies. Finally, *P. trifoliata* was used as focal genome and synteny diversity was calculated using the adapted scripts from the authors of the method. The genome assembly for *C. australis* was reverse complemented for the pseudomolecules “GWHBQDX00000001”, “GWHBQDX00000003”, “GWHBQDX00000006”, and “GWHBQDX00000007”, so they were on the same orientation than the remaining 8 genome assemblies in those pseudomolecules.

### Determination of synteny in pairwise comparison between NLR clusters

For the nine genome assemblies used on the synteny diversity analysis, syntenic pairs of NLR clusters were defined as those clusters from genome 1 that overlap at least 15% of their length with syntenic alignment(s) that simultaneously covered 15% of the length of another NLR cluster in genome 2.

## Acknowledgements

We thank Dr. Tracy Kahn for curating the Givaudan Citrus Variety Collection at UCR and providing access to ‘Washington’ navel for sampling. This research was supported by USDA-NIFA Awards # 2020-70029-33202 and 2023-70029-41305 to D.K.S; USDA-NIFA Award # 2019-70016-29796 to G.C.; Citrus Research Board Award # 5200-171 D.K.S. T.R.B is a fellow in the Plants-3D NSF National Research Traineeship Program (DBI-1922642). E.A.D is a UC Mexus Postdoctoral Fellow. Computations were performed using the computer clusters and data storage resources of the HPCC, which were funded by grants from NSF (MRI-2215705, MRI-1429826) and NIH (1S10OD016290-01A1).

## Author contributions

E.A.D, T.R.B, S.P.T, G.C, and D.K.S. conceived and planned the research. E.A.D, T.R.B, S.P.T, D.M.P performed analysis and contributed to the interpretation of the results. E.A.D, T.R.B, D.K.S drafted the initial manuscript and all authors provided critical feedback and edited the manuscript. G.C and D.K.S provided funding.

## Data availability

RNA-sequencing reads and Pacbio HiFi reads for *C. sinensis* ‘Washington navel’ have been deposited in NCBI under the Bioproject PRJNA989187. Genome assembly for *C. sinensis* ‘Washington navel’ has been deposited in NCBI under the Bioprojects PRJNA970396 and PRJNA970399 for haplotypes A and B respectively.

The raw sequence data are available in the Sequence Read Archive under NCBI BioProject PRJNA951227, PRJNA942352 for *C. limon* ‘Limoneira 8A’ and PRJNA951226, PRJNA942351 for *C. reticulata* ‘Sun Chu Sha Kat’. Genome assembly for *C. limon* ‘Limoneira 8A’ has been deposited in NCBI under accession JAWIMB000000000 and JAWIMC000000000. Genome assembly for *C. reticulata* ‘Sun Chu Sha Kat’ has been deposited in NCBI under accession JAWHPW000000000 and JAWHPX000000000.

## Conflict of interest statement

The authors declare no conflicts of interest.

## Supplementary Figure Legends

**Figure S1.** Genome quality assessment with Merqury spectrum plot for *C. limon* ‘Limoneira 8A’, *C. reticulata*‘Sun Chu Sha Kat’, and *C. sinensis* ‘Washington navel’ for evaluating K-mer completeness. K-mer multiplicity refers to the number of times a k-mers is found in sequencing reads. Colors correspond to the number of times a k-mer is found in the assembly. Nomenclature: *C. limon* ‘Limoneira 8A’ (*Citrus limon*), *C. reticulata* ‘Sun Chu Sha Kat’ (SCSK), *C. sinensis* ‘Washington navel’ (WN).

**Figure S2.** Comparison of LAIs for Citrus genomes; *C. sinensis* WN_HapB, *C. sinensis* WN_HapA, *C. reticulata* SCSK_Hap2, *C. reticulata* SCSK_Hap1, *C. limon* PB449_Hap2, *C. limon* PB449_Hap1, *C. ichangensis*, *C. sinensis* DVS, *C. sinensis* HZAU, *C. sinensis*, *C. limon*, *C. clementina*, *C. reticulata* T3, *C. reticulata*, *C. australis*, *P. trifoliata*. The red line indicates the average of LAI score across all genomes. The star denotes the genome assemblies generated in this work. Nomenclature: *C. limon* ‘Limoneira 8A’ (*C. limon* PB449), *C. reticulata* ‘Sun Chu Sha Kat’ (SCSK), *C. sinensis* ‘Washington navel’ (WN), *C. sinensis* ‘Valencia’ (DVS and HZAU).

**Figure S3.** Distribution of tandem repeats across the *C. sinensis* ‘Washington navel’ (A, B), *C. limon*‘Limoneira 8A’ (C, D), and *C. reticulata* ‘Sun Chu Sha Kat’ (E, F) genomes. Circos plots of tandem repeats identified by TRASH. Shading is colored according to repeat length, red corresponds to 181-bp repeats, blue to 460-bp repeats, green to 156-bp repeats, and purple to 336-bp repeats.

**Figure S4.** Phylogenetic tree for the consensus sequences from 181-bp and one 178-bp tandem repeats in *C. limon* ‘Limoneira 8A’, *C. reticulata* ‘Sun Chu Sha Kat’, and *C. sinensis* ‘Washington navel’ genomes. The tree was built using the Neighbor-Joining method. The percentage of replicate trees in which the associated sequences clustered together in the bootstrap test (1000 replicates) are shown above the branches (only those above 50). The evolutionary distances were computed using the Jukes-Cantor. The analysis involved 92 repeat sequences. All positions with less than 90% site coverage were eliminated, i.e., fewer than 10% alignment gaps, missing data, and ambiguous bases were allowed at any position. There were a total of 163 positions in the final dataset. Nomenclature: *C. limon* ‘Limoneira 8A’ (Limoneira 8A), *C. reticulata* ‘Sun Chu Sha Kat’ (SCSK), *C. sinensis* ‘Washington navel’ (WN), Sour Orange (SO).

**Figure S5.** A comparison of WGDs in *C. limon* ‘Limoneira 8A’, *C. reticulata* ‘Sun Chu Sha Kat’, and *C. sinensis* ‘Washington navel’ with Citrus species and other plant species. Nomenclature: *C. limon* ‘Limoneira 8A’ (*Citrus limon*), *C. reticulata* ‘Sun Chu Sha Kat’ (SCSK), *C. sinensis* ‘Washington navel’ (WN), Whole-genome duplication (WGD).

**Figure S6.** Distribution of NLR genes in chromosomes of A) *C. limon* ‘Limoneira 8A’, B) *C. reticulata* ‘Sun Chu Sha Kat’, and C) *C. sinensis* ‘Washington navel’.

**Figure S7**. Distribution of the intergenic distance calculated for all the genes in citrus (pink) or only NLRs (aquamarine).

**Figure S8.** Relationship between number of physical clusters of NLRs and inter-NLR distance in citrus (left) and number of NLRs found in the cluster with the most NLRs (right). NLRs were clustered based on the physical distance between adjacent NLRs in each genome, allowing up to 50 kb between adjacent NLRs to be considered part of the same cluster.

**Figure S9.** Synteny diversity analysis for nine chromosome-level genome assemblies. Yellow bars highlight the physical localization of NLR clusters in *P. trifoliata*. Scores were calculated for 200-Kb windows with a 100-Kb step. The black line represents the scores along the genome. The dotted lines point out synteny diversity scores of 0.417 (blue) and 0.722 (brown). *P. trifoliata* was used as the focal genome.

**Figure S10.** The distribution of synteny diversity score for permuted genomic regions. Permuted regions were the same length as observed NLR clusters in *P. trifoliata*. The distribution represents the synteny diversity mean scores for a Monte Carlo test with 10,000 permutations of 80 random genomic locations each time. The vertical red line highlights the mean synteny diversity score observed for the 80 NLR clusters in *P. trifoliata*.

**Figure S11.** Distribution of the number of genes in a cluster (left) and the cluster length (right) for clusters containing genes in multiple orthogroups (OGs, blue) or not (brown). Multi-orthogroup clusters of NLR tended to have more genes and be longer than single-orthogroup clusters (Gene number: W = 192100, p= 4.58^-14^; Cluster length: W = 201976, p=7.94^-10^). Significance was calculated using a Mann-Whitney U test.

**Figure S12.** Distance to the closest transposable element (TE) for any gene, an NLRs, or an NLR with integrated domain (ID) in *C. sinensis* ‘Washington navel’. A Welch’s ANOVA was used to identify significant differences between the means of the groups. A Games-Howel post hoc test was used to identify significant in which groups the significant differences were, represented by a distinct letter on top of the box plot for each category (*p* <= 0.05).

**Figure S13.** Proportion of secondary alignments within an NLR cluster with respect to the primary alignment in citrus. A lower proportion suggests that the cluster contains mainly non-duplicated sequences. A score of 1 represents that the cumulative length of secondary alignments is equal to the length of the NLR cluster.

**Figure S14.** Dotplot of self-alignment of cluster 89 in *C. sinensis* ‘Washington navel’ haplotype B. Duplicated regions within the cluster can be seen as secondary alignments (off-diagonal alignments). Only alignments of at least one kilobase are shown.

**Figure S15.** Diagram showing the distribution of NLRs for a single cluster across nine chromosome-level genome assemblies used to calculate synteny diversity. Arrows indicate NLRs, whereas their color represents the orthogroup classification. For each genome assembly the scale represents 50 Kb. Green, blue and yellow bars shown for *C. sinensis* WN haplotype B and *C. limon* haplotype sour orange represent duplicated regions. Nomenclature: *C. limon* ‘Limoneira 8A’ (*C. limon*), *C. reticulata* ‘Sun Chu Sha Kat’ (SCSK), *C. sinensis*‘Washington navel’ (WN).

**Figure S16.** Phylogenetic tree for NLRs contained in the set of clusters syntenic to cluster 89 in *C. sinensis*‘Washington navel’ haplotype B. The tree was built using the Neighbor-Joining method. The percentage of replicate trees in which the associated sequences clustered together in the bootstrap test (1000 replicates) are shown above the branches (only those above 50). The evolutionary distances were computed using the JTT matrix-based method. The analysis involved 210 amino acid sequences. All positions with less than 90% site coverage were eliminated, i.e., fewer than 10% alignment gaps, missing data, and ambiguous bases were allowed at any position. There were a total of 859 positions in the final dataset. Nomenclature: *C. limon*‘Limoneira 8A’ (LL), *C. reticulata* ‘Sun Chu Sha Kat’ (SCSK), *C. sinensis* ‘Washington navel’ (WN), *C. clementina* (Clementina), *C. australis* (Australis), sour orange (orange), *P. trifoliata* (Trifoliata). Figure S17. Phylogeny and diagram of the physical position of genes duplicates in cluster 89 in *C. sinensis* ‘Washington navel’ haplotype B. The black and purple lines connect the phylogenetic tree with the physical position within the cluster for a set of duplicated NLRs. The colors in NLR names represent if they belong to the most frequent orthogroup (green) or not (brown). Tree was rooted using the ZAR1 NLR in *Arabidopsis thaliana*, and the evolutionary distances were computed using the JTT matrix-based method. The analysis involved 36 amino acid sequences.

**Figure S17**. Phylogeny and diagram of the physical position of genes duplicates in cluster 89 in *C. sinensis* ‘Washington navel’ haplotype B. The black and purple lines connect the phylogenetic tree with the physical position within the cluster for a set of duplicated NLRs. The colors in NLR names represent if they belong to the most frequent orthogroup (green) or not (brown). Tree was rooted using the ZAR1 NLR in *Arabidopsis thaliana*, and the evolutionary distances were computed using the JTT matrix-based method. The analysis involved 36 amino acid sequences.

**Figure S18.** Schematic representation of the sequences observed in *C. maxima* ‘Majiayou’, ‘Fairchild’ mandarin, a hypothetical haplotype from a *C. maxima* or *C. maxima* hybrid, and *C. limon* ‘Limoneira 8A’ haplotype sour orange for the duplicated region in Cluster 80. A hypothetical recombination between haplotypes to create the current cluster 80 is shown in the blue x symbol. Boxes with numbers represent NLRs, whereas boxes with the word “RT” represent retrotransposable elements, boxes with the word “E4” represent the E4 region. The blue box represents a retrotransposable element that was found in *C. maxima-*derived haplotypes but missing in *C. reticulata*-derived haplotypes.

**Figure S19.** Nucleotide alignment from a section of the “E4” region showing evidence of unequal recombination for *C. limon* ‘Limoneira 8A’ haplotype sour orange, *C. maxima* ‘Huazhouyou-tomentosa’ (HZYT), and ‘Fairchild’ mandarin. The 25-bp region (light blue) is also found after the 55-bp indel (light green).

## Notes

### Competing Interest Statement

The authors have declared no competing interest.

