## Supplemental Figures for "The spectrum of diversity of nucleotide-binding leucine-rich repeat (NLR) genes in citrus and its relatives"

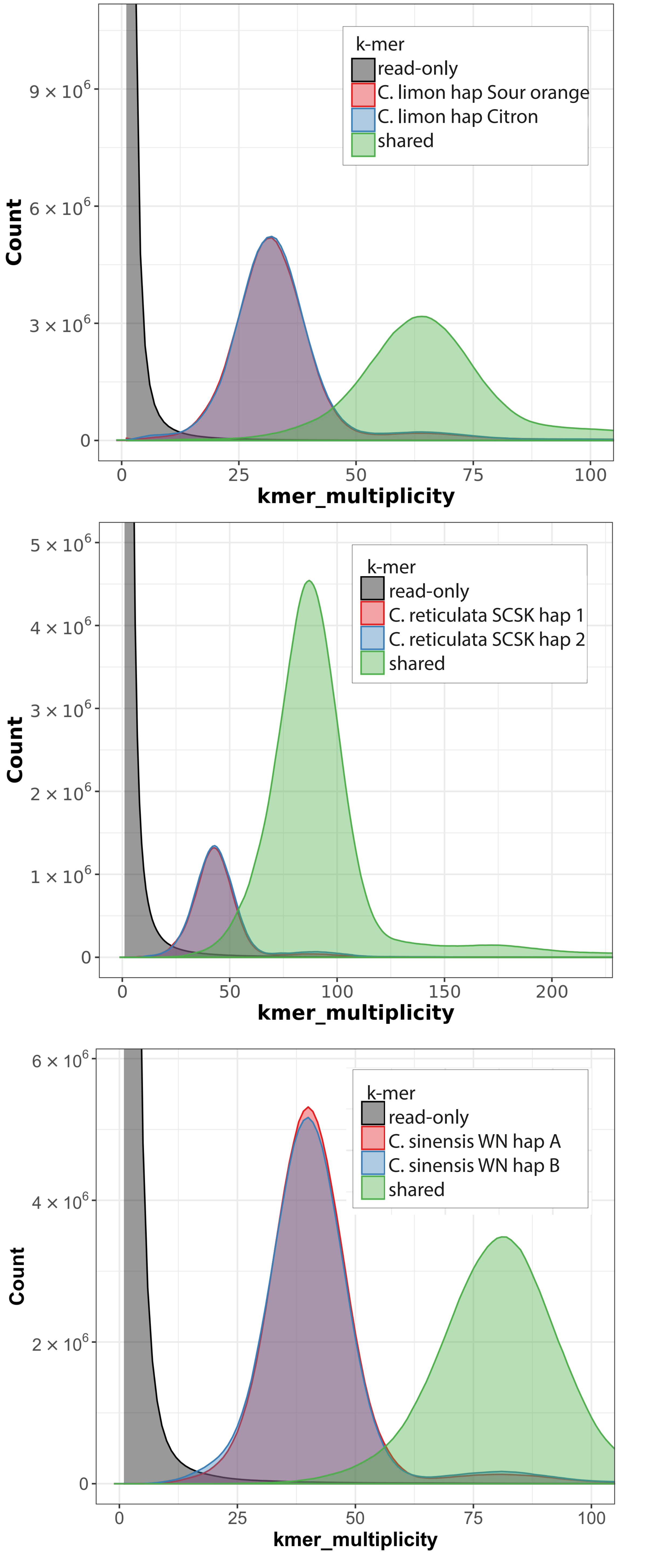

k-mer
read-only
C. limon hap Sour orange
C. limon hap Citron
shared
Count
kmer_multiplicity
0
25
50
75
100
9 × 10^6
6 × 10^6
3 × 10^6
k-mer
read-only
C. reticulata SCSK hap 1
C. reticulata SCSK hap 2
shared
Count
kmer_multiplicity
0
50
100
150
200
5 × 10^6
4 × 10^6
3 × 10^6
2 × 10^6
1 × 10^6
k-mer
read-only
C. sinensis WN hap A
C. sinensis WN hap B
shared
Count
kmer_multiplicity
0
25
50
75
100
6 × 10^6
4 × 10^6
2 × 10^6

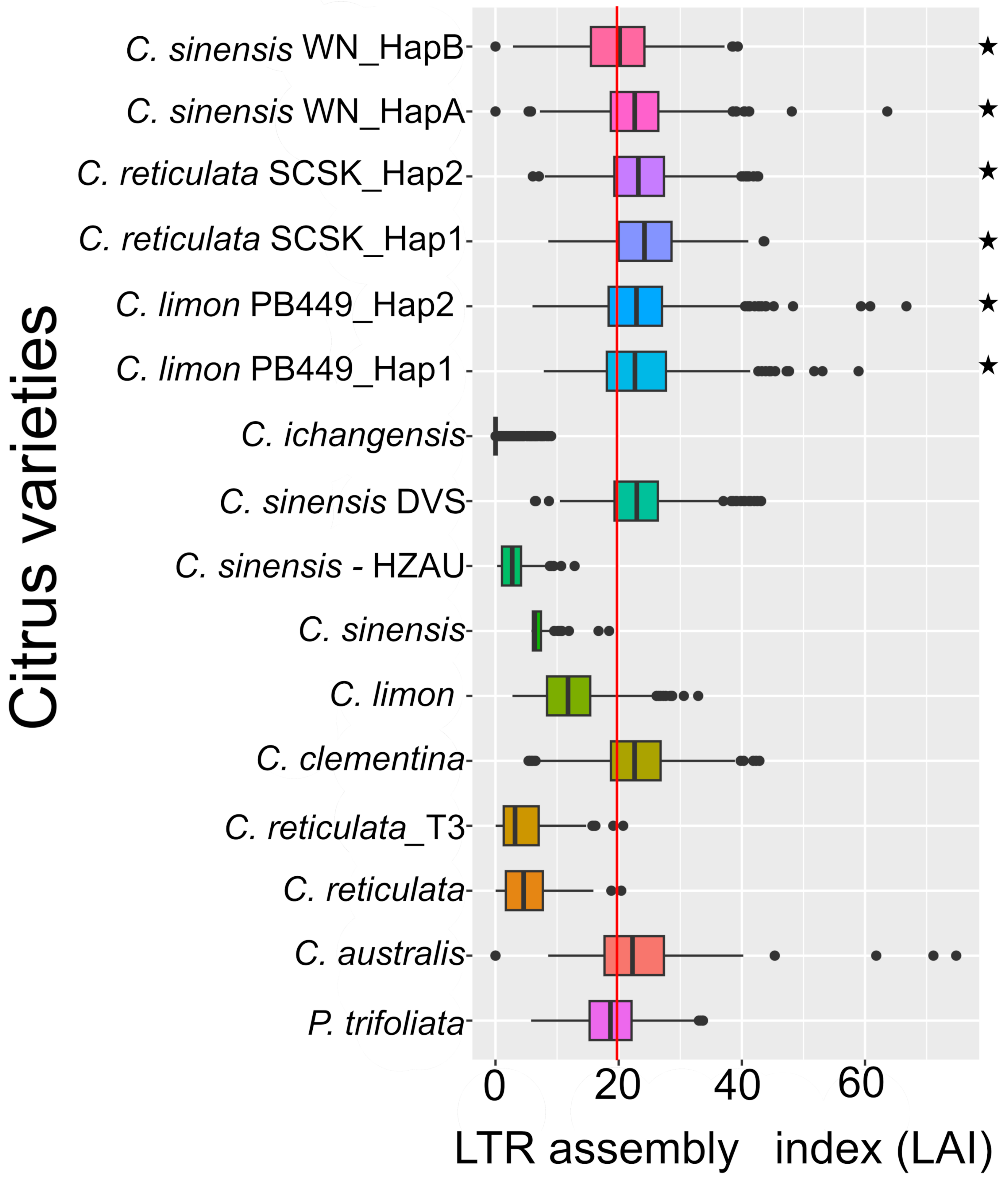
C. sinensis WN_HapB
C. sinensis WN_HapA
C. reticulata SCSK_Hap2
C. reticulata SCSK_Hap1
C. limon PB449_Hap2
C. limon PB449_Hap1
C. ichangensis
C. sinensis DVS
C. sinensis - HZAU
C. sinensis
C. limon
C. clementina
C. reticulata_T3
C. reticulata
C. australis
P. trifoliata
Citrus varieties
0
20
40
60
LTR assembly index (LAI)

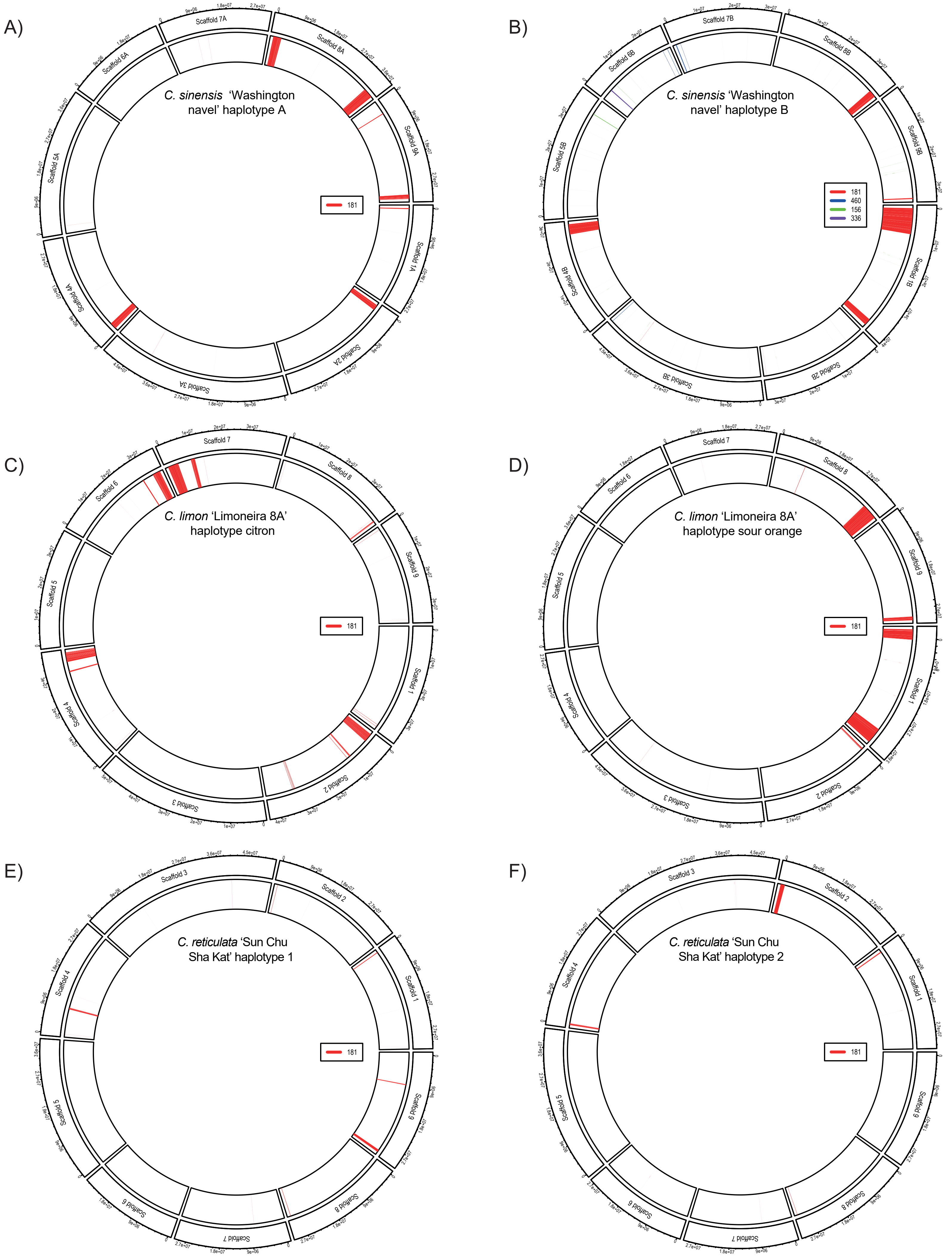
A)
C. sinensis 'Washington navel' haplotype A
181
Scaffold 1A
Scaffold 2A
Scaffold 3A
Scaffold 4A
Scaffold 5A
Scaffold 6A
Scaffold 7A
Scaffold 8A
Scaffold 9A
B)
C. sinensis 'Washington navel' haplotype B
181
460
156
336
Scaffold 1B
Scaffold 2B
Scaffold 3B
Scaffold 4B
Scaffold 5B
Scaffold 6B
Scaffold 7B
Scaffold 8B
Scaffold 9B
C)
C. limon 'Limoneira 8A' haplotype citron
181
Scaffold 1
Scaffold 2
Scaffold 3
Scaffold 4
Scaffold 5
Scaffold 6
Scaffold 7
Scaffold 8
Scaffold 9
D)
C. limon 'Limoneira 8A' haplotype sour orange
181
Scaffold 1
Scaffold 2
Scaffold 3
Scaffold 4
Scaffold 5
Scaffold 6
Scaffold 7
Scaffold 8
Scaffold 9
E)
C. reticulata 'Sun Chu Sha Kat' haplotype 1
181
Scaffold 1
Scaffold 2
Scaffold 3
Scaffold 4
Scaffold 5
Scaffold 6
Scaffold 7
Scaffold 8
Scaffold 9
F)
C. reticulata 'Sun Chu Sha Kat' haplotype 2
181
Scaffold 1
Scaffold 2
Scaffold 3
Scaffold 4
Scaffold 5
Scaffold 6
Scaffold 7
Scaffold 8
Scaffold 9

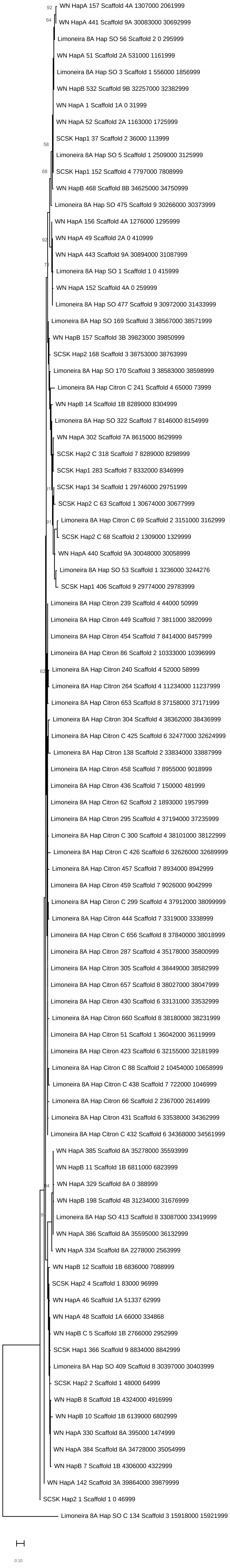
92
WN HapA 157 Scaffold 4A 1307000 2061999
64
WN HapA 441 Scaffold 9A 30083000 30692999
Limoneira 8A Hap SO 56 Scaffold 2 0 295999
WN HapA 51 Scaffold 2A 531000 1161999
Limoneira 8A Hap SO 3 Scaffold 1 556000 1856999
WN HapB 532 Scaffold 9B 32257000 32382999
WN HapA 1 Scaffold 1A 0 31999
WN HapA 52 Scaffold 2A 1163000 1725999
SCSK Hap1 37 Scaffold 2 36000 113999
58
Limoneira 8A Hap SO 5 Scaffold 1 2509000 3125999
68
SCSK Hap1 152 Scaffold 4 7797000 7808999
WN HapB 468 Scaffold 8B 34625000 34750999
Limoneira 8A Hap SO 475 Scaffold 9 30266000 30373999
WN HapA 156 Scaffold 4A 1276000 1295999
92
WN HapA 49 Scaffold 2A 0 410999
WN HapA 443 Scaffold 9A 30894000 31087999
77
Limoneira 8A Hap SO 1 Scaffold 1 0 415999
WN HapA 152 Scaffold 4A 0 259999
Limoneira 8A Hap SO 477 Scaffold 9 30972000 31433999
Limoneira 8A Hap SO 169 Scaffold 3 38567000 38571999
WN HapB 157 Scaffold 3B 39823000 39850999
SCSK Hap2 168 Scaffold 3 38753000 38763999
Limoneira 8A Hap SO 170 Scaffold 3 38583000 38598999
Limoneira 8A Hap Citron C 241 Scaffold 4 65000 73999
WN HapB 14 Scaffold 1B 8289000 8304999
Limoneira 8A Hap SO 322 Scaffold 7 8146000 8154999
WN HapA 302 Scaffold 7A 8615000 8629999
SCSK Hap2 C 318 Scaffold 7 8289000 8298999
SCSK Hap1 283 Scaffold 7 8332000 8346999
91
SCSK Hap1 34 Scaffold 1 29746000 29751999
SCSK Hap2 C 63 Scaffold 1 30674000 30677999
91
Limoneira 8A Hap Citron C 69 Scaffold 2 3151000 3162999
SCSK Hap2 C 68 Scaffold 2 1309000 1329999
WN HapA 440 Scaffold 9A 30048000 30058999
Limoneira 8A Hap SO 53 Scaffold 1 3236000 3244276
SCSK Hap1 406 Scaffold 9 29774000 29783999
Limoneira 8A Hap Citron 239 Scaffold 4 44000 50999
Limoneira 8A Hap Citron 449 Scaffold 7 3811000 3820999
Limoneira 8A Hap Citron 454 Scaffold 7 8414000 8457999
Limoneira 8A Hap Citron 86 Scaffold 2 10333000 10396999
62
Limoneira 8A Hap Citron 240 Scaffold 4 52000 58999
Limoneira 8A Hap Citron 264 Scaffold 4 11234000 11237999
Limoneira 8A Hap Citron 653 Scaffold 8 37158000 37171999
Limoneira 8A Hap Citron 304 Scaffold 4 38362000 38436999
Limoneira 8A Hap Citron C 425 Scaffold 6 32477000 32624999
Limoneira 8A Hap Citron 138 Scaffold 2 33834000 33887999
Limoneira 8A Hap Citron 458 Scaffold 7 8955000 9018999
Limoneira 8A Hap Citron 436 Scaffold 7 150000 481999
Limoneira 8A Hap Citron 62 Scaffold 2 1893000 1957999
Limoneira 8A Hap Citron 295 Scaffold 4 37194000 37235999
Limoneira 8A Hap Citron C 300 Scaffold 4 38101000 38122999
Limoneira 8A Hap Citron C 426 Scaffold 6 32626000 32689999
Limoneira 8A Hap Citron 457 Scaffold 7 8934000 8942999
Limoneira 8A Hap Citron 459 Scaffold 7 9026000 9042999
Limoneira 8A Hap Citron C 299 Scaffold 4 37912000 38099999
Limoneira 8A Hap Citron 444 Scaffold 7 3319000 3338999
Limoneira 8A Hap Citron C 656 Scaffold 8 37840000 38018999
Limoneira 8A Hap Citron 287 Scaffold 4 35178000 35800999
Limoneira 8A Hap Citron 305 Scaffold 4 38449000 38582999
Limoneira 8A Hap Citron 657 Scaffold 8 38027000 38047999
Limoneira 8A Hap Citron 430 Scaffold 6 33131000 33532999
Limoneira 8A Hap Citron 660 Scaffold 8 38180000 38231999
Limoneira 8A Hap Citron 51 Scaffold 1 36042000 36119999
Limoneira 8A Hap Citron 423 Scaffold 6 32155000 32181999
Limoneira 8A Hap Citron C 88 Scaffold 2 10454000 10658999
Limoneira 8A Hap Citron C 438 Scaffold 7 722000 1046999
Limoneira 8A Hap Citron 66 Scaffold 2 2367000 2614999
Limoneira 8A Hap Citron 431 Scaffold 6 33538000 34362999
Limoneira 8A Hap Citron C 432 Scaffold 6 34368000 34561999
WN HapA 385 Scaffold 8A 35278000 35593999
WN HapB 11 Scaffold 1B 6811000 6823999
64
WN HapA 329 Scaffold 8A 0 388999
WN HapB 198 Scaffold 4B 31234000 31676999
97
Limoneira 8A Hap SO 413 Scaffold 8 33087000 33419999
WN HapA 386 Scaffold 8A 35595000 36132999
WN HapA 334 Scaffold 8A 2278000 2563999
WN HapB 12 Scaffold 1B 6836000 7088999
SCSK Hap2 4 Scaffold 1 83000 96999
WN HapA 46 Scaffold 1A 51337 62999
WN HapA 48 Scaffold 1A 66000 334868
WN HapB C 5 Scaffold 1B 2766000 2952999
SCSK Hap1 366 Scaffold 9 8834000 8842999
Limoneira 8A Hap SO 409 Scaffold 8 30397000 30403999
SCSK Hap2 2 Scaffold 1 48000 64999
WN HapB 8 Scaffold 1B 4324000 4916999
WN HapB 10 Scaffold 1B 6139000 6802999
WN HapA 330 Scaffold 8A 395000 1474999
WN HapA 384 Scaffold 8A 34728000 35054999
WN HapB 7 Scaffold 1B 4306000 4322999
WN HapA 142 Scaffold 3A 39864000 39879999
SCSK Hap2 1 Scaffold 1 0 46999
Limoneira 8A Hap SO C 134 Scaffold 3 15918000 15921999
0.10

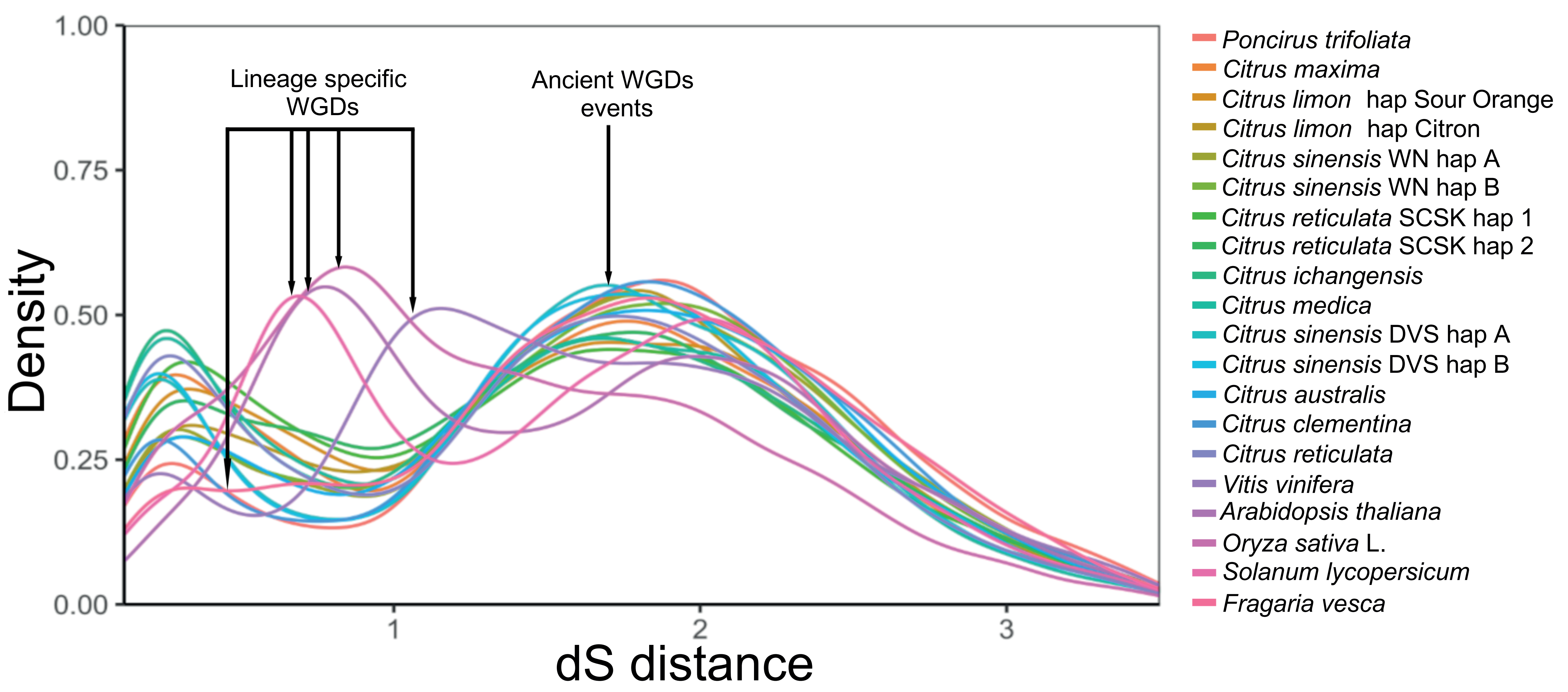

Lineage specific WGDs
Ancient WGDs events
Density
dS distance
1.00
0.75
0.50
0.25
0.00
1
2
3
Poncirus trifoliata
Citrus maxima
Citrus limon hap Sour Orange
Citrus limon hap Citron
Citrus sinensis WN hap A
Citrus sinensis WN hap B
Citrus reticulata SCSK hap 1
Citrus reticulata SCSK hap 2
Citrus ichangensis
Citrus medica
Citrus sinensis DVS hap A
Citrus sinensis DVS hap B
Citrus australis
Citrus clementina
Citrus reticulata
Vitis vinifera
Arabidopsis thaliana
Oryza sativa L.
Solanum lycopersicum
Fragaria vesca

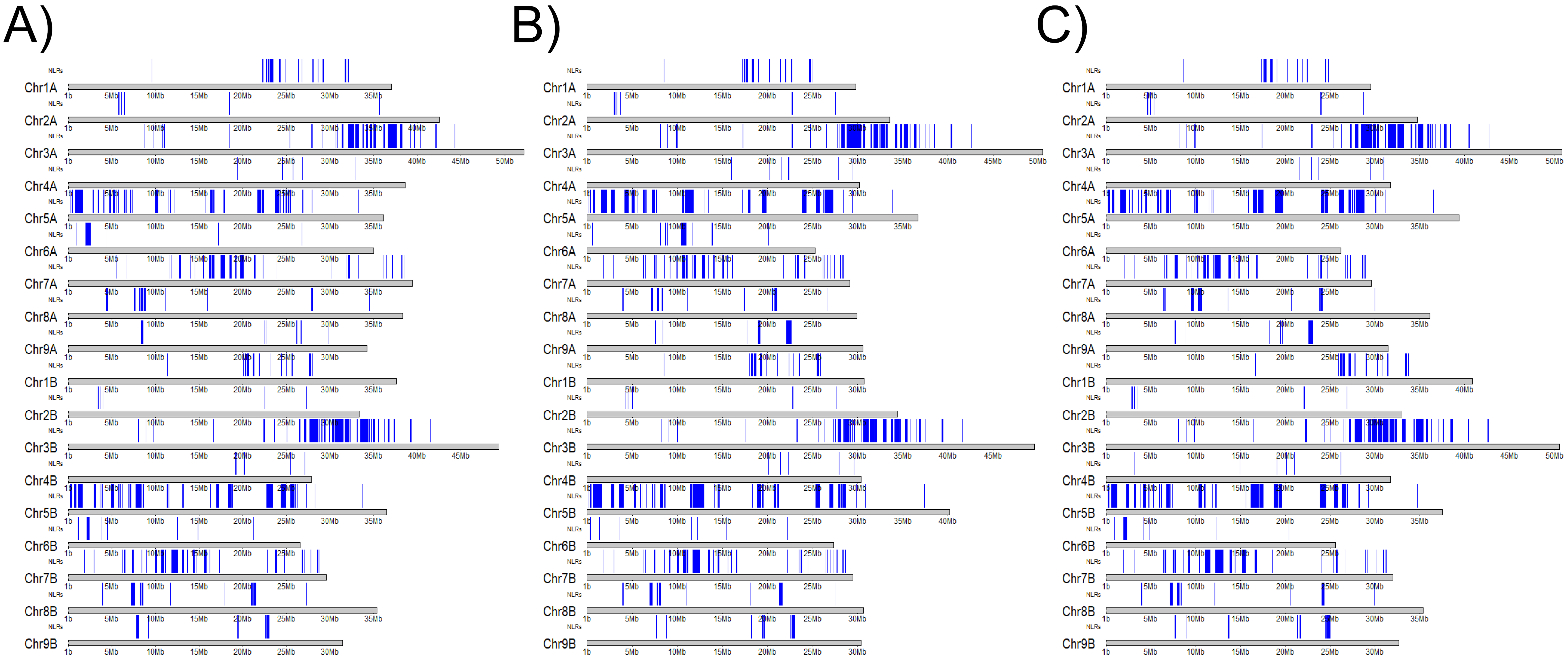

A)
B)
C)
NLRs
Chr1A
Chr2A
Chr3A
Chr4A
Chr5A
Chr6A
Chr7A
Chr8A
Chr9A
Chr1B
Chr2B
Chr3B
Chr4B
Chr5B
Chr6B
Chr7B
Chr8B
Chr9B
1b
5Mb
10Mb
15Mb
20Mb
25Mb
30Mb
35Mb
40Mb
45Mb
50Mb

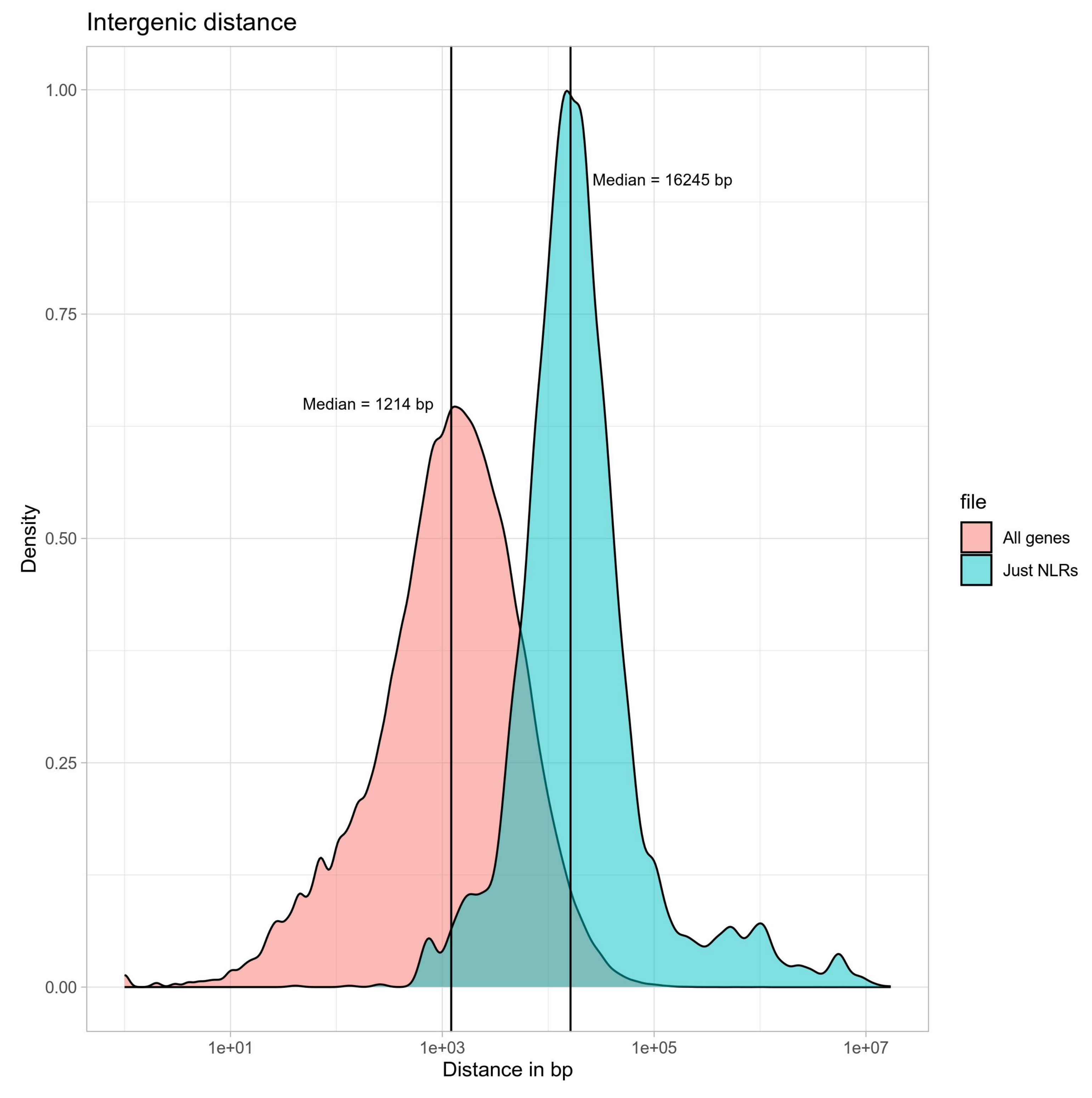
Intergenic distance
Median = 16245 bp
Median = 1214 bp
Density
1.00
0.75
0.50
0.25
0.00
1e+01
1e+03
1e+05
1e+07
Distance in bp
file
All genes
Just NLRs

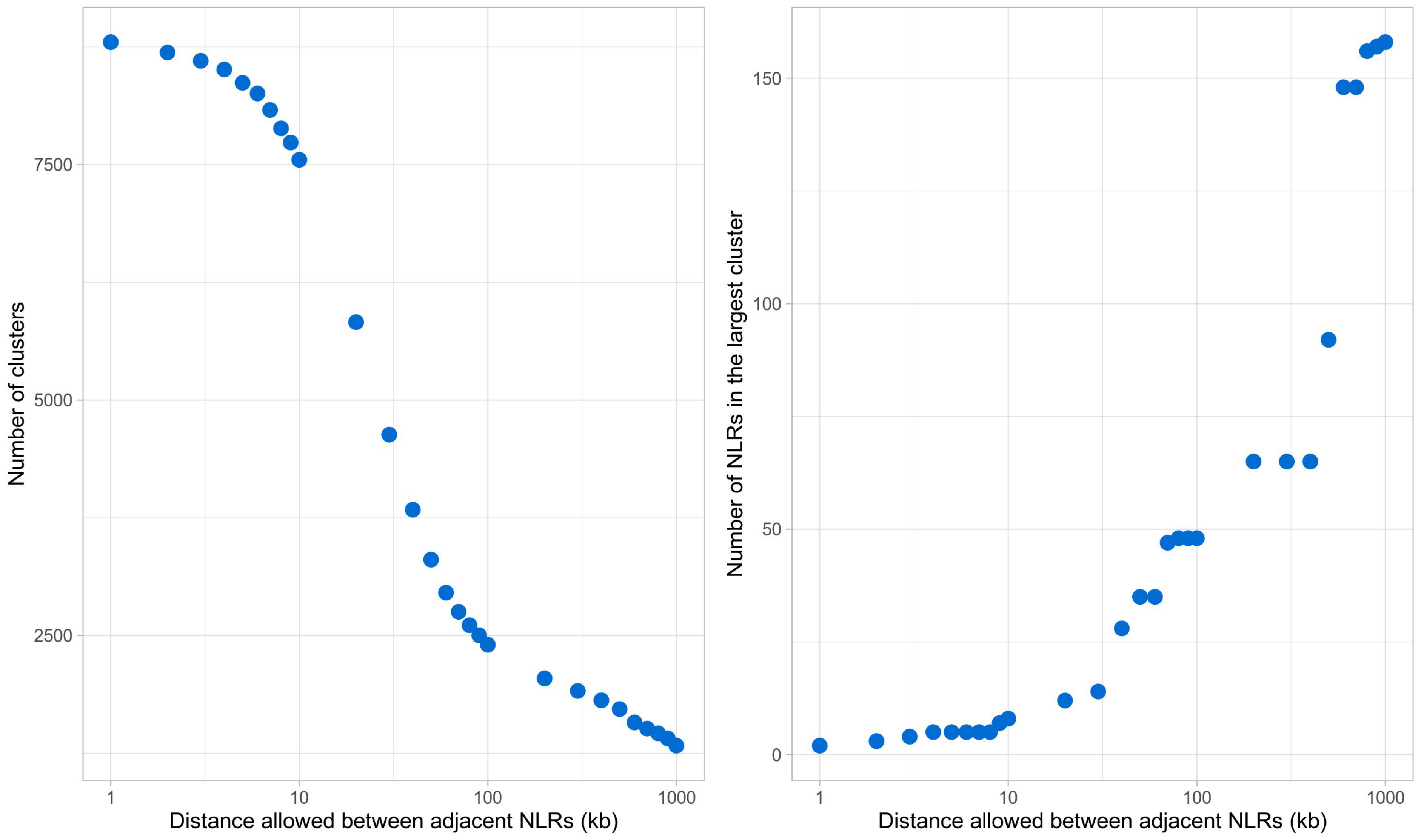

Number of clusters
7500
5000
2500
1
10
100
1000
Distance allowed between adjacent NLRs (kb)
Number of NLRs in the largest cluster
150
100
50
0
1
10
100
1000
Distance allowed between adjacent NLRs (kb)

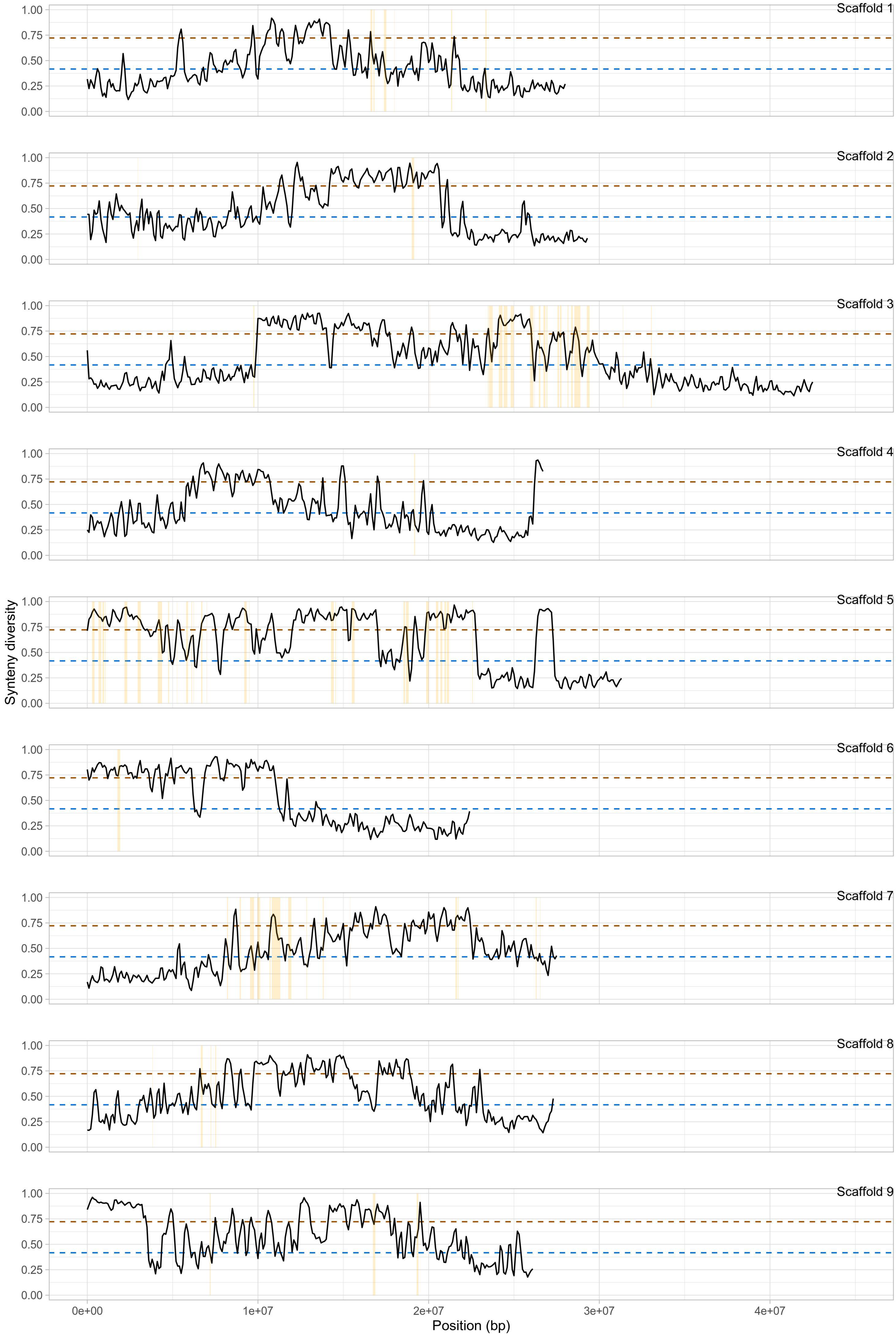
Scaffold 1
Scaffold 2
Scaffold 3
Scaffold 4
Scaffold 5
Scaffold 6
Scaffold 7
Scaffold 8
Scaffold 9
1.00
0.75
0.50
0.25
0.00
Synteny diversity
0e+00
1e+07
2e+07
3e+07
4e+07
Position (bp)

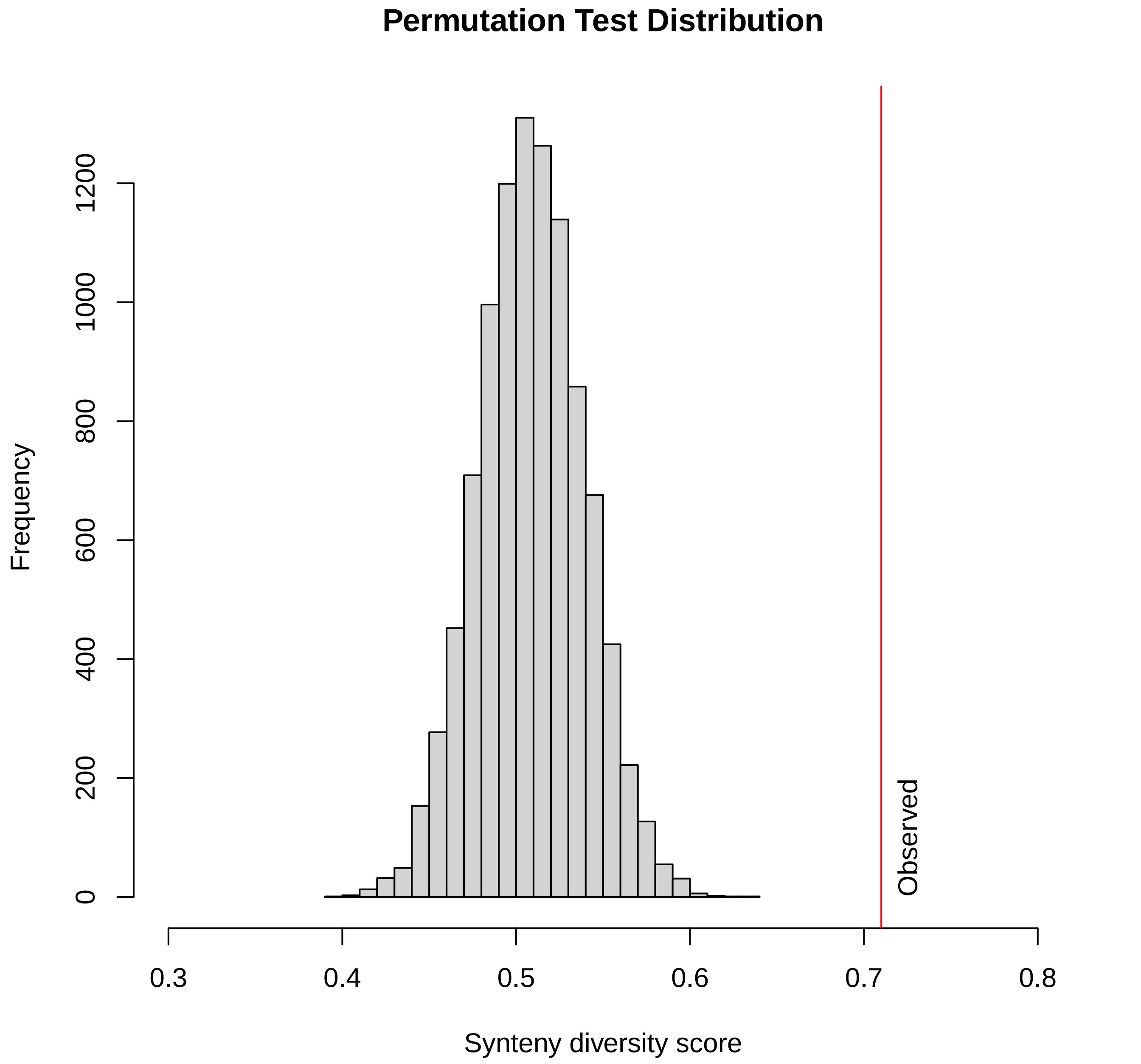

Permutation Test Distribution
Frequency
0
200
400
600
800
1000
1200
Observed
0.3
0.4
0.5
0.6
0.7
0.8
Synteny diversity score

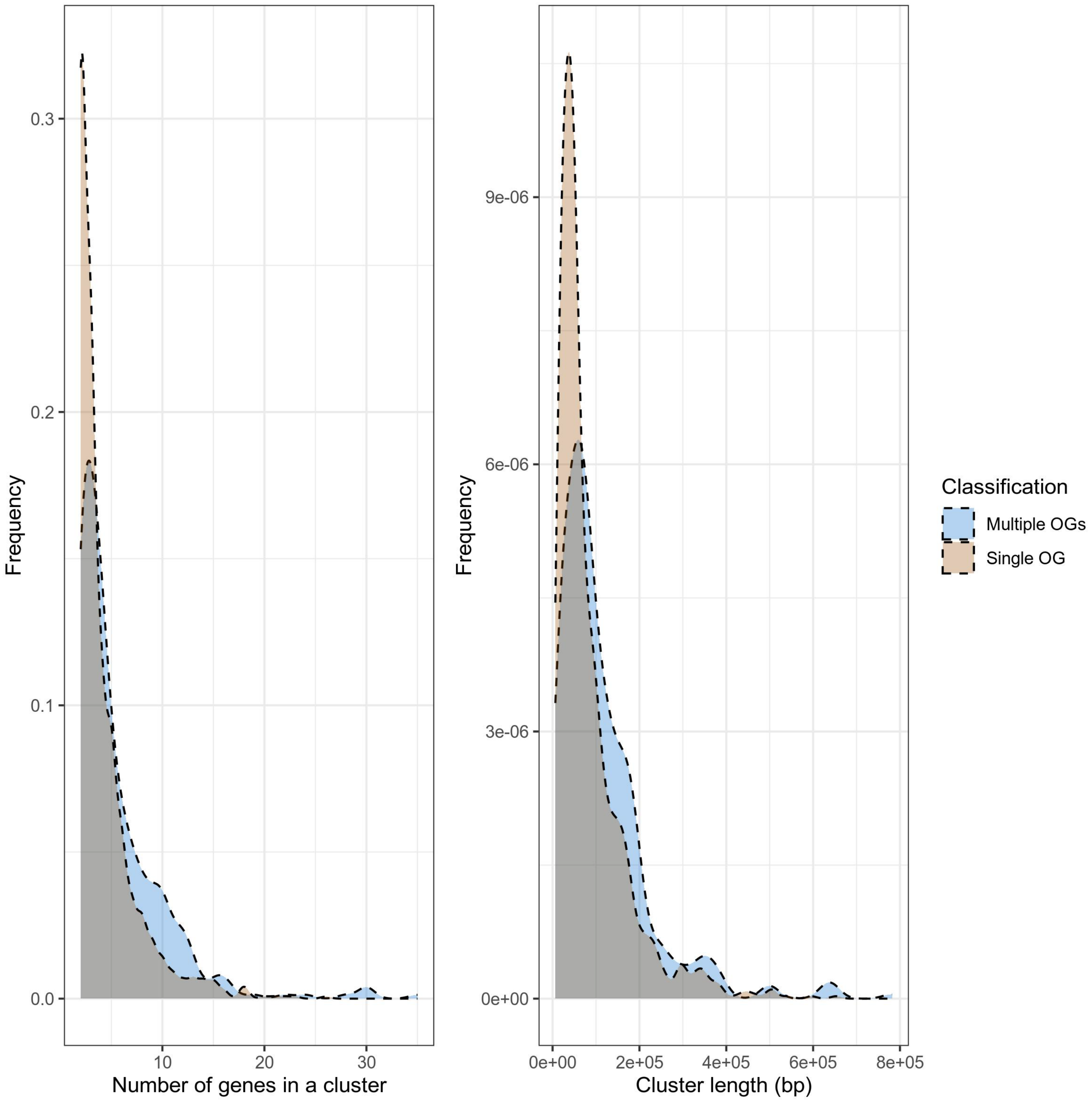

Frequency
0.3
0.2
0.1
0.0
10
20
30
Number of genes in a cluster
Frequency
9e-06
6e-06
3e-06
0e+00
0e+00
2e+05
4e+05
6e+05
8e+05
Cluster length (bp)
Classification
Multiple OGs
Single OG

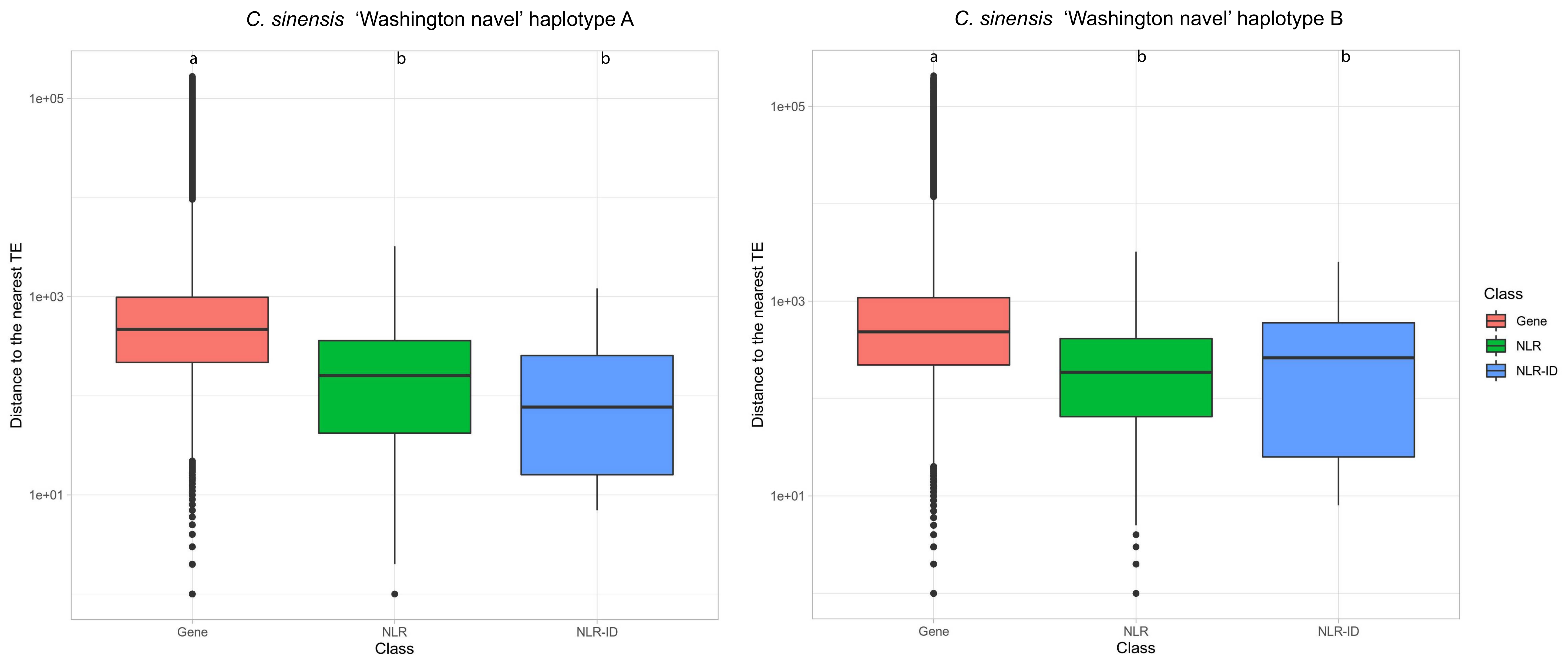
C. sinensis 'Washington navel' haplotype A
C. sinensis 'Washington navel' haplotype B
a
b
b
a
b
b
1e+05
1e+03
1e+01
Distance to the nearest TE
Gene
NLR
NLR-ID
Class
Gene
NLR
NLR-ID
NLR-ID

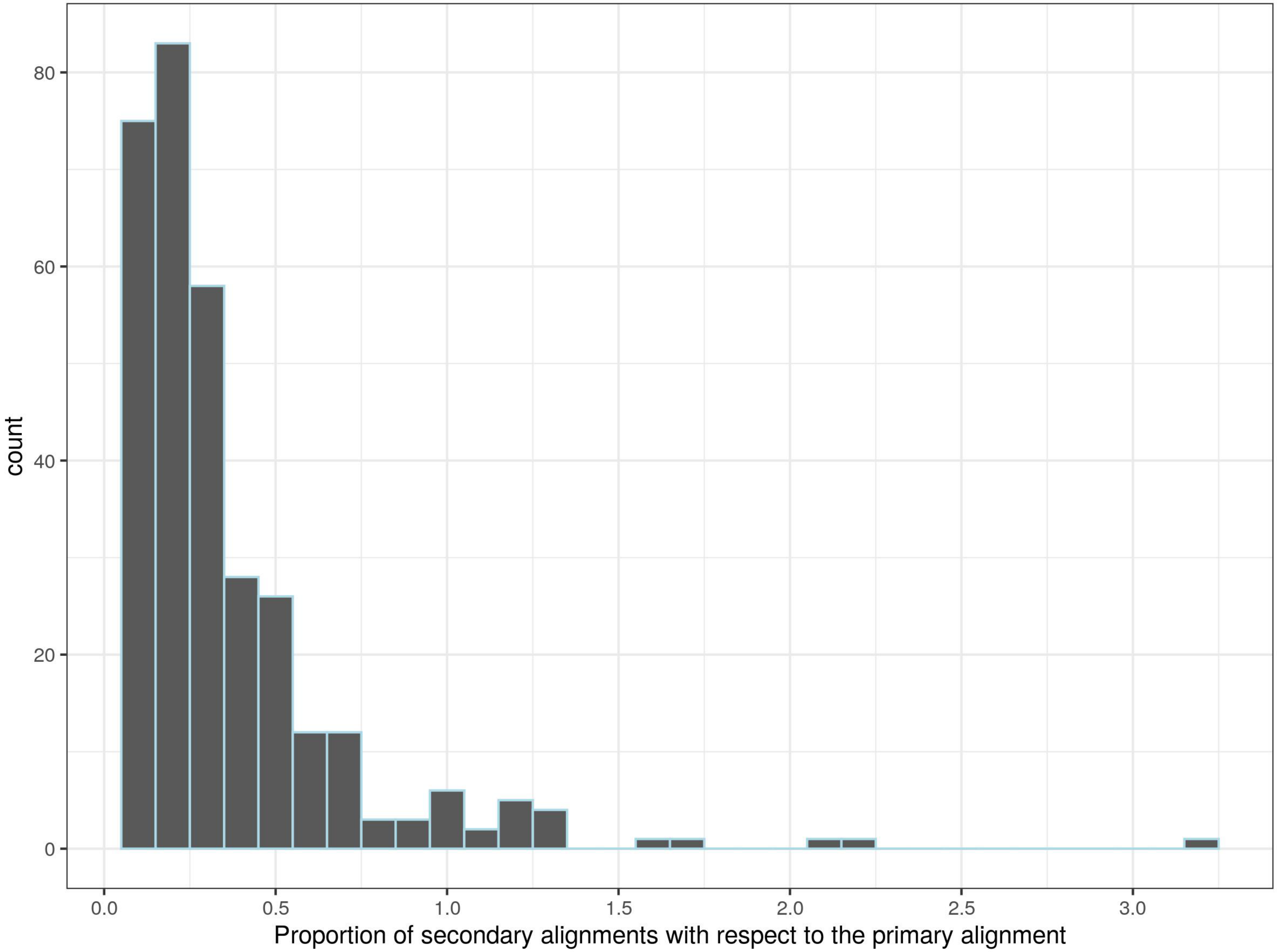

count
80
60
40
20
0
0.0
0.5
1.0
1.5
2.0
2.5
3.0
Proportion of secondary alignments with respect to the primary alignment

Post-filtering number of alignments: 651 minimum alignment length (-m): 1000
Post-filtering number of queries: 1 minimum query aggregate alignment length (-q): 3000

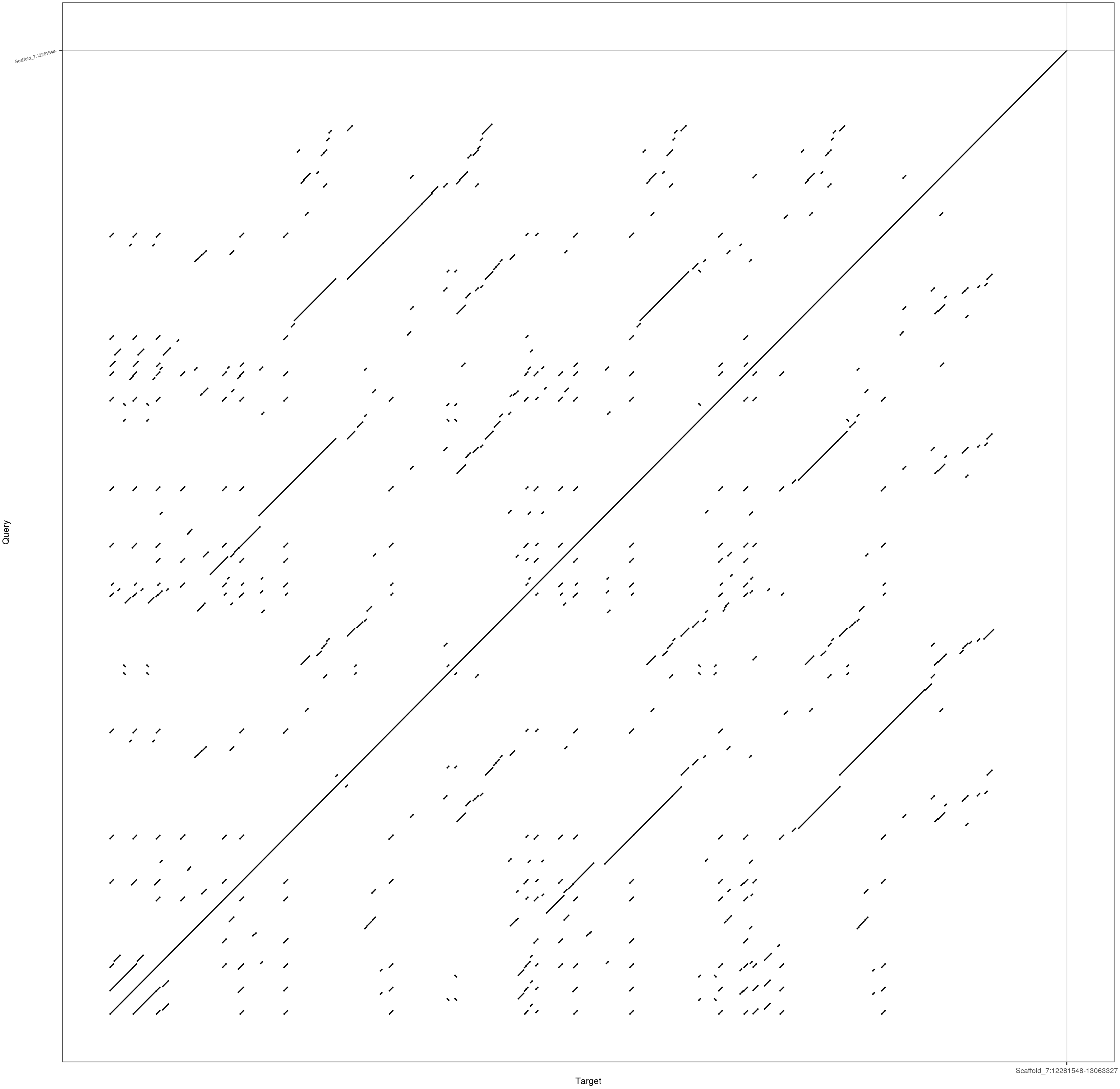

**Syntenic clusters to C65 in *P. trifoliata***

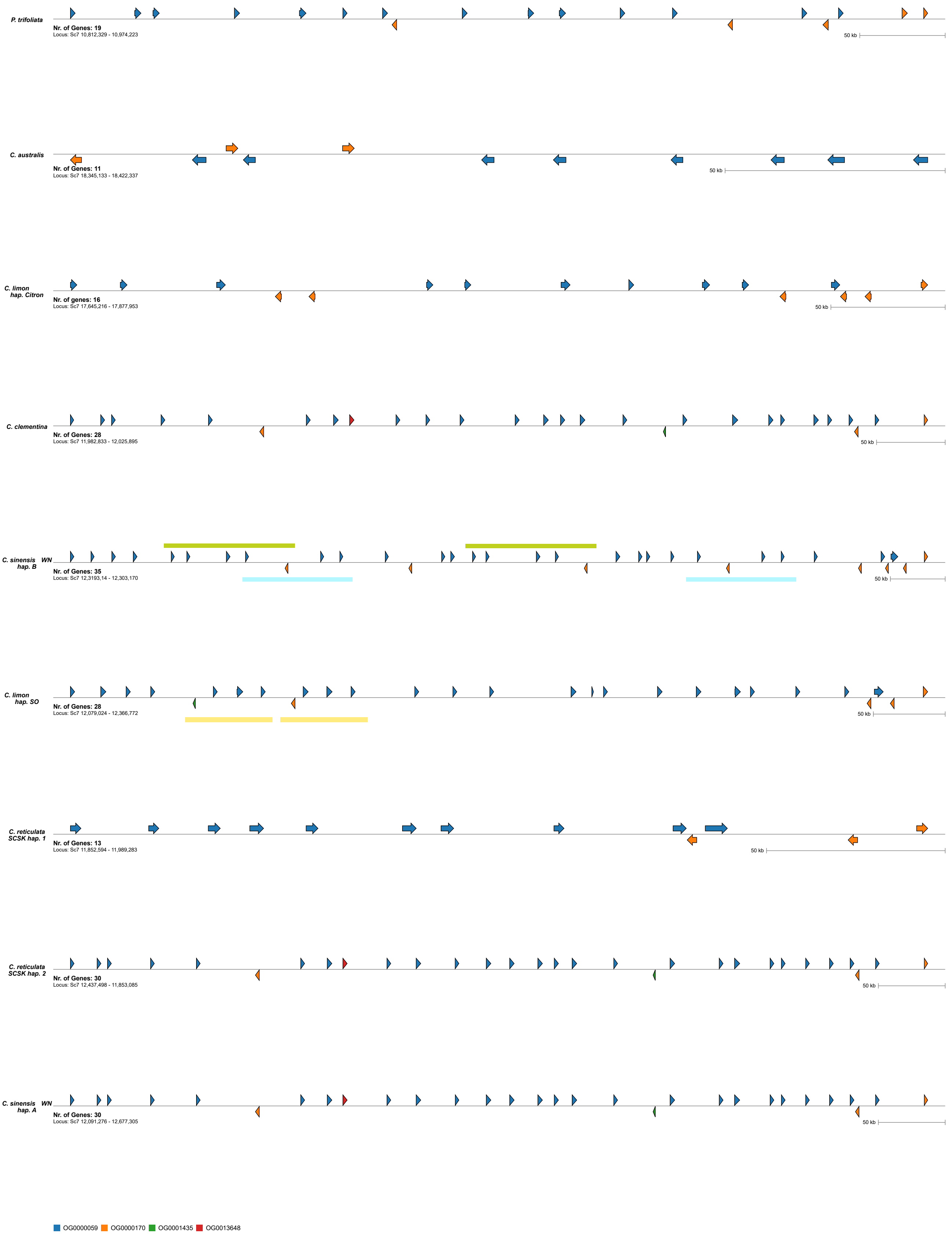

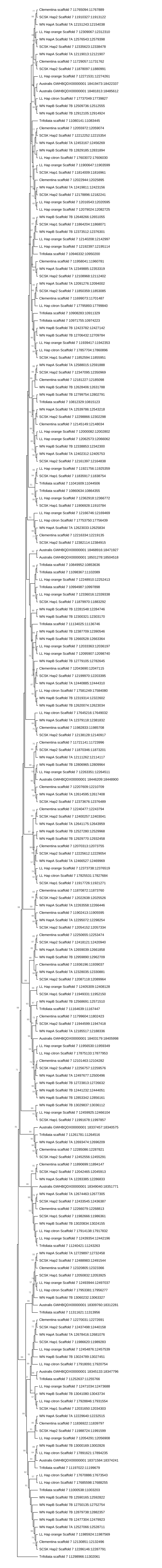

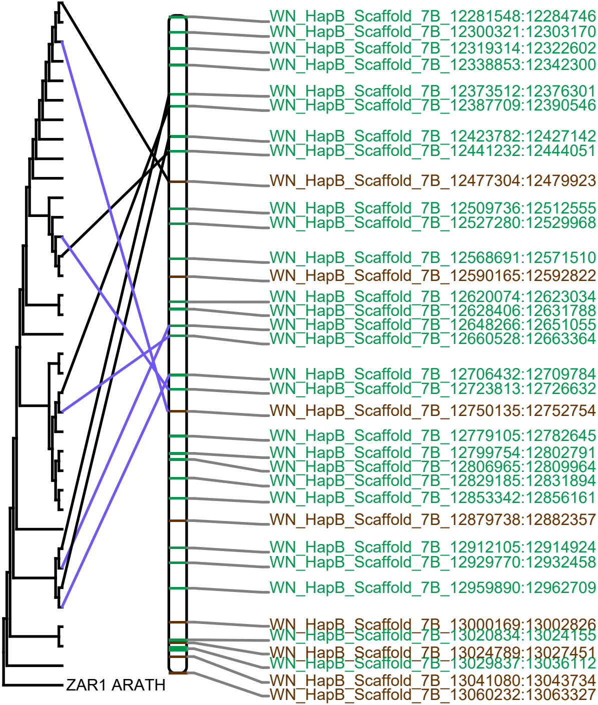
WN_HapB_Scaffold_7B_12281548:12284746
WN_HapB_Scaffold_7B_12300321:12303170
WN_HapB_Scaffold_7B_12319314:12322602
WN_HapB_Scaffold_7B_12338853:12342300
WN_HapB_Scaffold_7B_12373512:12376301
WN_HapB_Scaffold_7B_12387709:12390546
WN_HapB_Scaffold_7B_12423782:12427142
WN_HapB_Scaffold_7B_12441232:12444051
WN_HapB_Scaffold_7B_12477304:12479923
WN_HapB_Scaffold_7B_12509736:12512555
WN_HapB_Scaffold_7B_12527280:12529968
WN_HapB_Scaffold_7B_12568691:12571510
WN_HapB_Scaffold_7B_12590165:12592822
WN_HapB_Scaffold_7B_12620074:12623034
WN_HapB_Scaffold_7B_12628406:12631788
WN_HapB_Scaffold_7B_12648266:12651055
WN_HapB_Scaffold_7B_12660528:12663364
WN_HapB_Scaffold_7B_12706432:12709784
WN_HapB_Scaffold_7B_12723813:12726632
WN_HapB_Scaffold_7B_12750135:12752754
WN_HapB_Scaffold_7B_12779105:12782645
WN_HapB_Scaffold_7B_12799754:12802791
WN_HapB_Scaffold_7B_12806965:12809964
WN_HapB_Scaffold_7B_12829185:12831894
WN_HapB_Scaffold_7B_12853342:12856161
WN_HapB_Scaffold_7B_12879738:12882357
WN_HapB_Scaffold_7B_12912105:12914924
WN_HapB_Scaffold_7B_12929770:12932458
WN_HapB_Scaffold_7B_12959890:12962709
WN_HapB_Scaffold_7B_13000169:13002826
WN_HapB_Scaffold_7B_13020834:13024155
WN_HapB_Scaffold_7B_13024789:13027451
WN_HapB_Scaffold_7B_13029837:13036112
WN_HapB_Scaffold_7B_13041080:13043734
WN_HapB_Scaffold_7B_13060232:13063327
ZAR1 ARATH

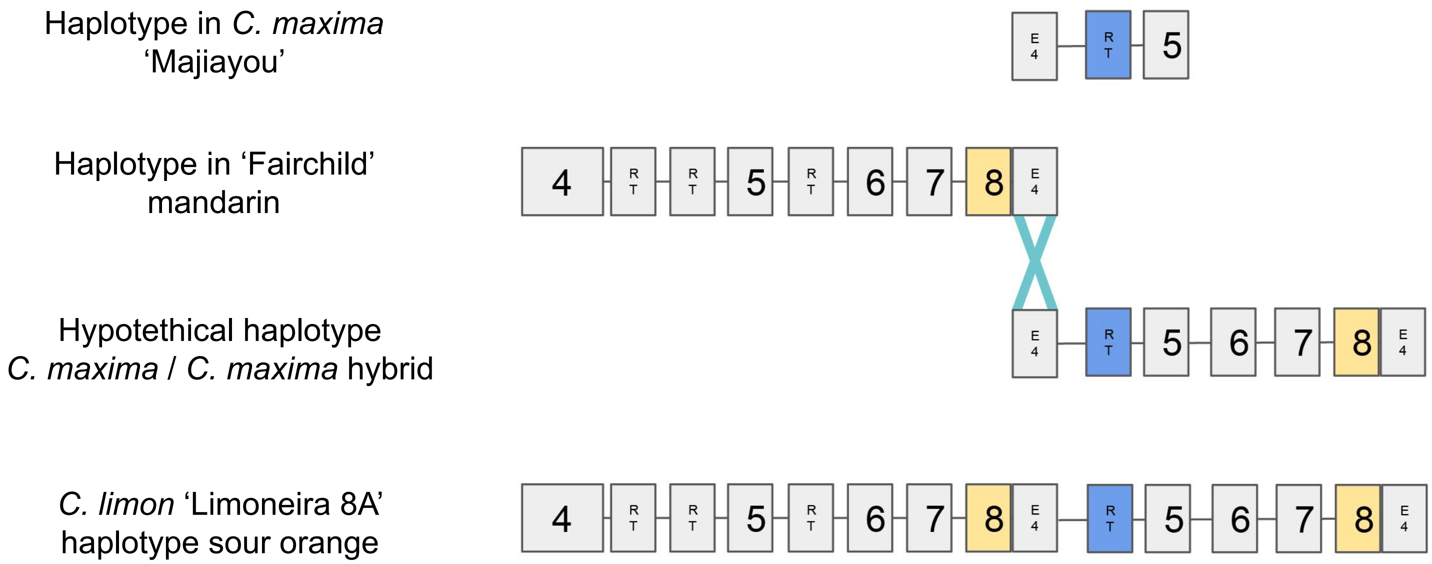
Haplotype in *C. maxima* 'Majiayou'
E 4
R T
5
Haplotype in 'Fairchild' mandarin
4
R T
R T
5
R T
6
7
8
E 4
Hypotethical haplotype *C. maxima* / *C. maxima* hybrid
E 4
R T
5
6
7
8
E 4
*C. limon* 'Limoneira 8A' haplotype sour orange
4
R T
R T
5
R T
6
7
8
E 4
R T
5
6
7
8
E 4

```
                                               4241
C. limon 'Limoneira 8A' hap SO - E4 Region 5'  TGAAAAATGT GTAGTTTTTG TTCCTTTTTT TCTTAAAAAT
C. maxima 'HZYT'                               TGAAAAATGT GTAGTTTTTG TTCCTTTTTT TCTTAAAAAT
Fairchild mandarin                             TGAAAAATGT GTAGTTTTTG TTCC-TTTTT TCTTAAAAAT
C. limon 'Limoneira 8A' hap SO - E4 Region 3'  TGAAAACTGT GTAGTTTTTG TTCC-TTTTT TCTTAAAAAT

                                               4281
C. limon 'Limoneira 8A' hap SO - E4 Region 5'  GGGAAAAAAG AATGTAA-TT TGTTTAAATT ACTGTTGTGA
C. maxima 'HZYT'                               GGGAAAAAAG AATGTAATTT TGTTTAAATT ACTGTTGTGA
Fairchild mandarin                             GGG-AAAAAG AATGTAATTT TGTTTAAATT ACTGTTGTGA
C. limon 'Limoneira 8A' hap SO - E4 Region 3'  GGG-AAAAAG AATGTAATTT TGTTTAAATT ACTGTTGTGA

                                               4321
C. limon 'Limoneira 8A' hap SO - E4 Region 5'  TAACAATTAT TATTTTATTG ATTAATTGGC AGAGA-----
C. maxima 'HZYT'                               TAACAATTAT TATTTTATTG ATTAATTGGC AGAGA-----
Fairchild mandarin                             TAACAATTAT TATTTTATTG ATTAATTGGC AGAGAATCGT
C. limon 'Limoneira 8A' hap SO - E4 Region 3'  TAACAATTAT TATTTTATTG ATTAATTGGC AGAGAATCGT

                                               4361
C. limon 'Limoneira 8A' hap SO - E4 Region 5'  ---------- ---------- ---------- ----------
C. maxima 'HZYT'                               ---------- ---------- ---------- ----------
Fairchild mandarin                             TATGGTTTCG TATGGTTGTT TTTTTTTTTT TTTCGTATGG
C. limon 'Limoneira 8A' hap SO - E4 Region 3'  TATGGTTTCG TATGGTTGTT TTTTTTTTTT TTTCGTATGG

                                               4401
C. limon 'Limoneira 8A' hap SO - E4 Region 5'  ---------- ATCGTTATGG TTTCGTATGG TTGTTAAAAC
C. maxima 'HZYT'                               ---------- ATCGTTATGG TTTCGTATGG TTGTTAAAAC
Fairchild mandarin                             TTGTTATGGT ATCGTTATGG TTTCGTATGG TTGTTAAAAC
C. limon 'Limoneira 8A' hap SO - E4 Region 3'  TTGTTATGGT ATCGTTATGG TTTCGTATGG TTGTTAAAAC

                                               4441
C. limon 'Limoneira 8A' hap SO - E4 Region 5'  ATTGGAAAGT CCACTACATC ATCCGTTAAT AACTTTCAAA
C. maxima 'HZYT'                               ATTGGAAAGT CCACTACATC ATCCGTTAAT AACTTTCAAA
Fairchild mandarin                             ATTGGAAAGT CCACTACATC ATCCGTTAAT AACTTTCAAA
C. limon 'Limoneira 8A' hap SO - E4 Region 3'  ATTGGAAAGT CCACTACATC ATCCGTTAAT AACTTTCAAA

                                               4481
C. limon 'Limoneira 8A' hap SO - E4 Region 5'  TTTATGGTTA ATCAATAGTT GAGATTTATT ATCATTAATG
C. maxima 'HZYT'                               TTTATGGTTA ATCAATAGTT GAGATTTATT ATCATTAATG
Fairchild mandarin                             TTTATGGTTA ATCAATAGTT GAGATTTATT ATCATTAAGG
C. limon 'Limoneira 8A' hap SO - E4 Region 3'  TTTATGGTTA ATCAATAGTT GAGATTTATT ATCATTAAGG
```
